# CXCR3-expressing metastasis-initiating cells induce and exploit a fibroblast niche in the lungs to fuel metastatic colonization

**DOI:** 10.1101/546952

**Authors:** Maren Pein, Jacob Insua-Rodríguez, Jasmin Meier, Tsunaki Hongu, Lena Wiedmann, Marieke A.G. Essers, Hans-Peter Sinn, Saskia Spaich, Marc Sütterlin, Andreas Schneeweiss, Andreas Trumpp, Thordur Oskarsson

**Affiliations:** Heidelberg Institute for Stem Cell Technology and Experimental Medicine (HI-STEM gGmbH), Im Neuenheimer Feld 280, 69120 Heidelberg, Germany; Division of Stem Cells and Cancer, German Cancer Research Center (DKFZ), Im Neuenheimer Feld 280, 69120 Heidelberg, Germany; Faculty of Biosciences, University of Heidelberg, Germany; Division of Stress-induced Activation of Hematopoietic Stem Cells, German Cancer Research Center (DKFZ), Im Neuenheimer Feld 280, 69120 Heidelberg, Germany; Institute of Pathology, University of Heidelberg, Im Neuenheimer Feld 224, 69120, Heidelberg, Germany; Department of Obstetrics and Gynaecology, University Medical Centre Mannheim, Heidelberg University, Theodor-Kutzer-Ufer 1-3, 68167, Mannheim, Germany; National Center for Tumor Diseases, Heidelberg University Hospital, German Cancer Research Center, Im Neuenheimer Feld 460, 69120 Heidelberg, Germany; DKFZ-ZMBH Alliance, 69120 Heidelberg, Germany; German Cancer Consortium (DKTK), 69120 Heidelberg, Germany

## Abstract

Metastatic colonization relies on interactions between disseminated cancer cells and the microenvironment in secondary organs. Here, we show that disseminated breast cancer cells evoke major phenotypic changes in lung fibroblasts to form a metastatic niche that supports malignant growth. Colonization of the lungs by cancer cells confers an inflammatory phenotype in associated fibroblasts, where IL-1α and IL-1β, secreted by breast cancer cells, induce *CXCL9* and *CXCL10* production in metastasis-associated fibroblasts via NF-κB signaling. These paracrine interactions fuel the growth of lung metastases. Notably, we find that the chemokine receptor CXCR3, that binds CXCL9/10, is specifically expressed in a small subset of breast cancer cells with stem/progenitor cell properties and high tumor-initiating ability when co-transplanted with fibroblasts. CXCR3-expressing cancer cells show high JNK signaling that drives IL-1α/β expression. Thus, CXCR3 marks a population of breast cancer cells that induces CXCL9/10 production in fibroblast, but can also respond to and benefit from these chemokines. Importantly, disruption of this intercellular JNK-IL-1-CXCL9/10-CXCR3 axis significantly reduces metastatic colonization in xenograft and syngeneic mouse models. These data mechanistically demonstrate an essential role for this molecular crosstalk between breast cancer cells and their fibroblast niche in the progression of metastasis.

## INTRODUCTION

Metastasis remains the primary threat to the lives of cancer patients with few effective therapeutic options available^1^. In breast cancer, metastases often occur years after the primary tumor has been diagnosed and resected. This indicates that the outgrowth of disseminated cancer cells towards clinically overt metastasis – metastatic colonization – is a rate-limiting step in the metastatic process. Indeed, despite high penetrance across cancer patient populations, successful metastatic colonization is inefficient at the cellular level in that disseminated cancer cells that invade secondary organs confront a suboptimal microenvironment and must cope with strong selective pressure^2^. Metastatic colonization is ultimately not only determined by genetic and epigenetic networks in cancer cells, but also by the microenvironment^3, 4^. Indeed, the non-transformed tumor microenvironment affects many aspects of cancer progression. A number of cells within the stroma, such as myeloid progenitor cells, macrophages, neutrophils, endothelial cells and fibroblasts have been implicated in tumor progression and metastasis. Disseminated cancer cells that successfully colonize secondary organs are able to not only withstand the repressive nature of resting microenvironment, but can also re-educate stromal cells to support malignant growth^5^. While the influence of the tumor microenvironment adds to the complexity of cancer initiation and progression, it likely also represents multiple opportunities for therapeutic intervention in metastatic cancer patients.

In cancer, the reactive microenvironment is recognized to have certain features of the microenvironment of healing wounds and regenerative tissues^6^. For example, fibroblasts, that form a heterogeneous group of mesenchymal cells that are commonly found within the connective tissues, play an essential role in tissue regeneration and wound healing^7, 8^. Upon tissue injury, fibroblasts alter their phenotype and acquire contractile properties and secrete large quantities of growth factors and extracellular matrix (ECM) proteins. Cancer associated fibroblasts (CAFs), that comprise significant portions of primary tumors, have received considerable attention in recent years and have been shown to regulate both tumor initiation and development^9^. In breast cancer, CAFs provide a cytokine and ECM milieu that promotes growth and progression of primary tumors^10–13^. Colonization of secondary organs by malignant cells requires modulation of the local microenvironment, leading to the formation of a metastatic niche that supports secondary tumor growth^5^. Whereas the understanding of CAF biology in primary malignancies is substantial and growing, the precise molecular function of stromal fibroblasts at metastatic sites and their effect on metastatic progression is poorly understood. This is particularly relevant when considering the dynamic stromal changes that occur during reprogramming of the microenvironment from early colonization to the growth of overt metastasis.

In this study, we explored the dynamic molecular interactions between disseminated breast cancer cells and fibroblasts during different stages of lung metastasis. We find that fibroblast number and phenotype change dramatically as metastatic nodules grow from micrometastases to macrometastases. Transcriptomic profiling of fibroblasts from metastatic lungs established by breast cancer cells with different metastatic potential reveals that highly metastatic cancer cells can induce early activation in fibroblasts characterized by a major increase in inflammatory and TGFβ signaling as well as proliferation. Two of the most highly induced genes in these metastasis-associated fibroblasts (MAFs) encode the inflammatory cytokines CXCL9 and CXCL10. We find that overexpression of these cytokines confers stem cell properties to breast cancer cells *in vitro* and promotes metastasis to the lungs in mouse models. Moreover, we find that a subset of breast cancer cells with high metastatic potential expresses the cell surface receptor CXCR3, that binds CXCL9 and CXCL10. Importantly, systemic treatment with an inhibitor of CXCR3 significantly reduces lung metastatic colonization of breast cancer cells in xenograft and syngeneic mouse models. Our data therefore reveals an important crosstalk between breast cancer cells and MAFs that promotes metastatic initiation and progression to overt lung metastasis.

## RESULTS

### Metastatic breast cancer cells induce early activation and inflammatory signaling in stromal lung fibroblasts

To investigate evolution of fibroblasts during metastatic colonization, we established experimental lung metastases by injecting MDA-MB-231 (MDA) human breast cancer cells or the highly metastatic derivative MDA-MB-231-LM2 (MDA-LM2) cells^14^ intravenously into immunocompromised mice. At one week post injection (when lungs harbor primarily micrometastases) and at three weeks post injection (when macrometastases are prominent and widespread), we isolated fibroblasts using fluorescence-activated cell sorting (FACS) (Fig. 1a-1c). Considering the different capacity of MDA and MDA-LM2 cells to grow metastasis in lungs, the experimental approach was designed to address the qualitative difference between MDA and MDA-LM2 associated fibroblasts in metastasis at each time point. Taking advantage of the heterogeneity within both populations, we selected individual mice for analysis that harbored comparable MDA or MDA-LM2 metastatic loads based on *in vivo* bioluminescence imaging (Supplementary Fig. 1a). Lung fibroblasts were isolated by FACS using two positive selection markers, PDGFRα and PDGFRβ, and a panel of negative selection markers (Fig. 1a and Supplementary Fig. 1b). Fibroblasts isolated from lungs with growing metastases were compared with fibroblasts from lungs of healthy, age-matched mice. Interestingly, we observed a substantial increase in the number of fibroblasts in lungs harboring macrometastases derived from both MDA and MDA-LM2 cell lines. In contrast, fibroblast numbers within micrometastases were comparable to those observed in healthy lungs (Fig. 1d). These data suggested that the fibroblast population in lung stroma expands extensively during metastatic colonization of breast cancer cells.

**Fig. 1.**
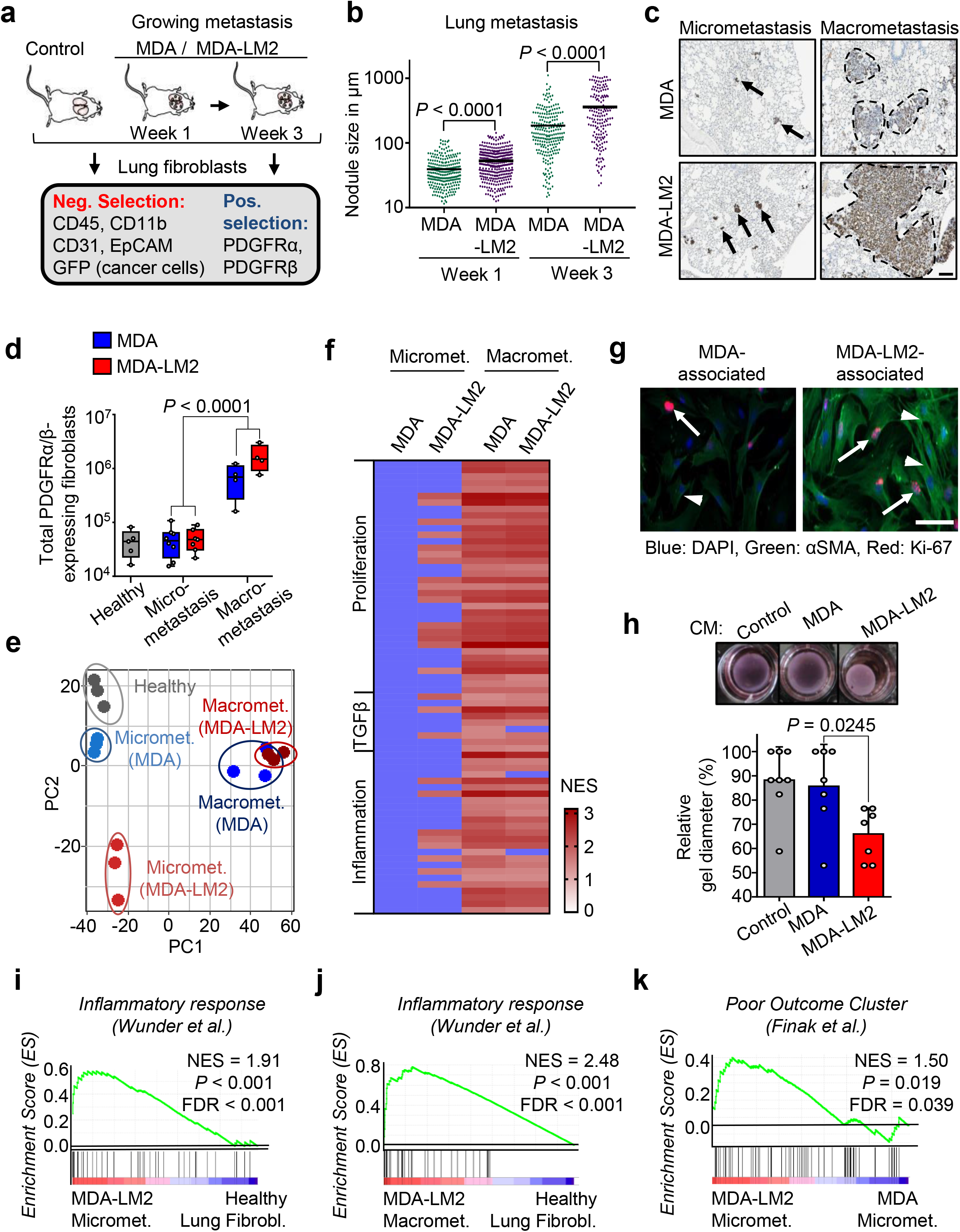
Metastatic breast cancer cells promote activation of lung fibroblasts during metastatic colonization that associates with poor prognosis. **a**, Schematic diagram of experimental setup for selection of fibroblasts from lungs of mice at different time points (week 1 and week 3) after intravenous injection of MDA-MB-231 (MDA) and MDA-MB-231-LM2 (MDA-LM2) breast cancer cells. **b**, Size of individual metastatic nodules at week 1 or week 3 after intravenous injection of MDA or MDA-LM2 breast cancer cells. Week 1, *n* = 257 metastatic nodules (MDA) and *n* = 330 metastatic nodules (MDA-LM2), from 6 mice each group. Week 3, *n* = 236 metastatic nodules (MDA) and *n* = 171 metastatic nodules (MDA-LM2), from 6 mice each group. **c**, Representative examples of MDA and MDA-LM2 micrometastases (from week 1) and macrometastases (from week 3) stained for human vimentin as a marker for cancer cells. Scale bar, 100 μM. Arrows indicate micrometastases and dotted lines indicate margins of macrometastases. **d**, Total number of fibroblasts from healthy lungs and metastatic lungs (MDA and MDA-LM2) at micrometastatic (week 1) and macrometastatic (week 3) stages. Healthy fibroblasts, *n* = 5 mice; fibroblasts from micrometastasis MDA, *n* = 8 mice and MDA-LM2, *n* = 7 mice; fibroblasts from macrometastasis (MDA or MDA-LM2), *n* = 4 mice each group. *P* value was determined by unpaired two-tailed t-test. **e**, Principal component (PC) analysis of transcriptome of fibroblasts isolated from metastatic lungs and healthy control lungs. **f**, Overview of GSEA using numerous gene signatures representing proliferation and TGFβ and inflammatory signaling. Heatmap shows normalized enrichment scores (NES) for signatures that were significantly changed, FDR < 0.1. Not significant changes compared to healthy lung fibroblasts are indicated by blue color. Gene Sets are provided in Supplementary Table 1. **g**, Immunocytochemical analysis of Ki-67 (red) and αSMA (green) expression in fibroblasts sorted from lungs 1 week after intravenous injection of MDA (left) or MDA-LM2 cells (right). Pictures are representative of 3 biological replicates. **h**, Contraction of collagen gels by NSG mouse lung fibroblasts stimulated with conditioned medium from MDA, MDA-LM2 and control medium. Control, *n* = 8, MDA or MDA-LM2, *n* = 7 for each group. *P* value was calculated by unpaired two-tailed t-test. **i-j**, Enrichment of an inflammatory response signature ^59^ in fibroblasts from MDA-LM2 micro- or macrometastasis compared to fibroblasts from healthy lungs. **k**, Enrichment of a poor outcome gene cluster ^60^ in fibroblasts isolated from MDA-LM2 compared to MDA micrometastases. For panels (i-k), NES, normalized enrichment score, FDR, false discovery rate. *P* values were determined by random permutation test.

To determine whether stromal lung fibroblasts phenotypically evolve as lung metastases progress, we performed transcriptomic analysis of purified fibroblasts. Principal component analysis (PCA) showed that biological replicates from each experimental group cluster together (Fig. 1e). Interestingly, fibroblasts from MDA-derived micrometastases, but not MDA-LM2-derived micrometastases, clustered close to healthy fibroblasts, whereas fibroblasts from macrometastases by both lines clustered together at a distance from healthy fibroblasts (Fig. 1e). Gene set enrichment analysis (GSEA) showed that, at the micrometastatic stage, MDA-LM2 breast cancer cells uniquely induced fibroblast activation based on early signs of proliferation and inflammation as well as TGFβ-signaling (Fig. 1f and Supplementary Table 1). At the macrometastatic stage, however, proliferation and inflammation signatures were strongly induced in MAFs by both breast cancer cell lines (Fig. 1f). Gene Ontology (GO) analysis revealed similar results in that the top 500 genes driving the PCA shift between MDA-LM2- and MDA-associated MAFs were notably involved in cell contraction, proliferation and inflammation (Supplementary Fig. 1c). In support of these findings, immunofluorescent staining of fibroblasts isolated from lungs harboring micrometastases showed that fibroblast proliferation and expression of alpha smooth muscle actin (αSMA), a marker for reactive fibroblasts, were increased in MDA-LM2-associated fibroblasts compared to MDA-associated fibroblasts (Fig. 1g and Supplementary Fig. 1d). Importantly, immunohistochemical staining of paraffin sections from human lung metastases from breast cancer patients revealed that 11/12 (92 %) samples exhibited αSMA-expressing fibroblasts (Supplementary Fig. 2a-c), whereas αSMA expression in healthy lungs was restricted to vessel linings (data not shown), indicating that activated MAFs are also implicated in human metastases. Within metastatic foci, αSMA-positive human fibroblasts were observed in direct contact with cancer cells (Supplementary Fig. 2a). Enhanced cell contractility in MDA-LM2-associated MAFs, suggested by GO analysis (Supplementary Fig. 1c), was functionally confirmed *in vitro*, as lung fibroblasts demonstrated a significant increase in collagen gel contraction upon stimulation with conditioned medium (CM) from MDA-LM2 cells compared to CM from MDA cells or control medium (Fig. 1h). Inflammatory response signatures were also observed in fibroblasts from MDA-LM2-derived micrometastases and were further enriched in macrometastases (Fig. 1f,i,J). Interestingly, fibroblasts associated with MDA-LM2 micrometastases showed a significant enrichment for genes comprising a stromal-derived “poor outcome” signature from breast cancer patients when compared to fibroblasts from lungs with MDA micrometastases (Fig. 1k). Moreover, this signature was further enriched in fibroblasts isolated from lungs with MDA and MDA-LM2 macrometastases (Supplementary Fig. 2d). These data support a model in which the phenotype of MAFs is influenced on one hand by the stage of metastatic progression and on the other the metastatic potential of associated cancer cells. Moreover, these data indicate that transcriptomic changes in MAFs are linked to poor outcome in breast cancer patients.

### *CXCL9* and *CXCL10* are induced in MAFs and promote lung metastasis in mouse models

Our findings led us to hypothesize that changes in stromal fibroblasts during metastatic colonization of the lungs may support the growth of metastasis. To address this, we aimed to identify genes expressed in MAFs that are involved in direct crosstalk with disseminated cancer cells and that are functionally relevant for metastatic growth in the lungs. Transcriptomic analysis of fibroblasts revealed that many genes encoding collagens, ECM glycoproteins or ECM modifying enzymes were markedly induced at macrometastatic stages (Supplementary Fig. 3a-c). Several of these genes have been shown to promote cancer and metastasis, such as *Tnc, Spp1, Fn1, Thbs2, Lox*, and *Serpinb2*^15–19^. Based on the link of early transcriptomic changes in MDA-LM2 associated fibroblasts to poor outcome (Fig. 1k), we reasoned that genes induced early in MDA-LM2 associated fibroblasts and further induced in macrometastases would be strong pro-metastatic candidates. Of the 115 genes that were induced in MAFs from MDA-LM2-derived micrometastases, 50 overlapped with genes expressed in MAFs from lungs harboring MDA- and MDA-LM2-derived macrometastases (Supplementary Fig. 4a,b), and this group comprised a number of genes encoding proteins that are secreted or membrane bound but exposed to the extracellular space. We prioritized these genes for further analysis and identified eight genes that represented candidates for a potential direct crosstalk between metastatic breast cancer cells and MAFs (Fig. 2a).

**Fig. 2.**
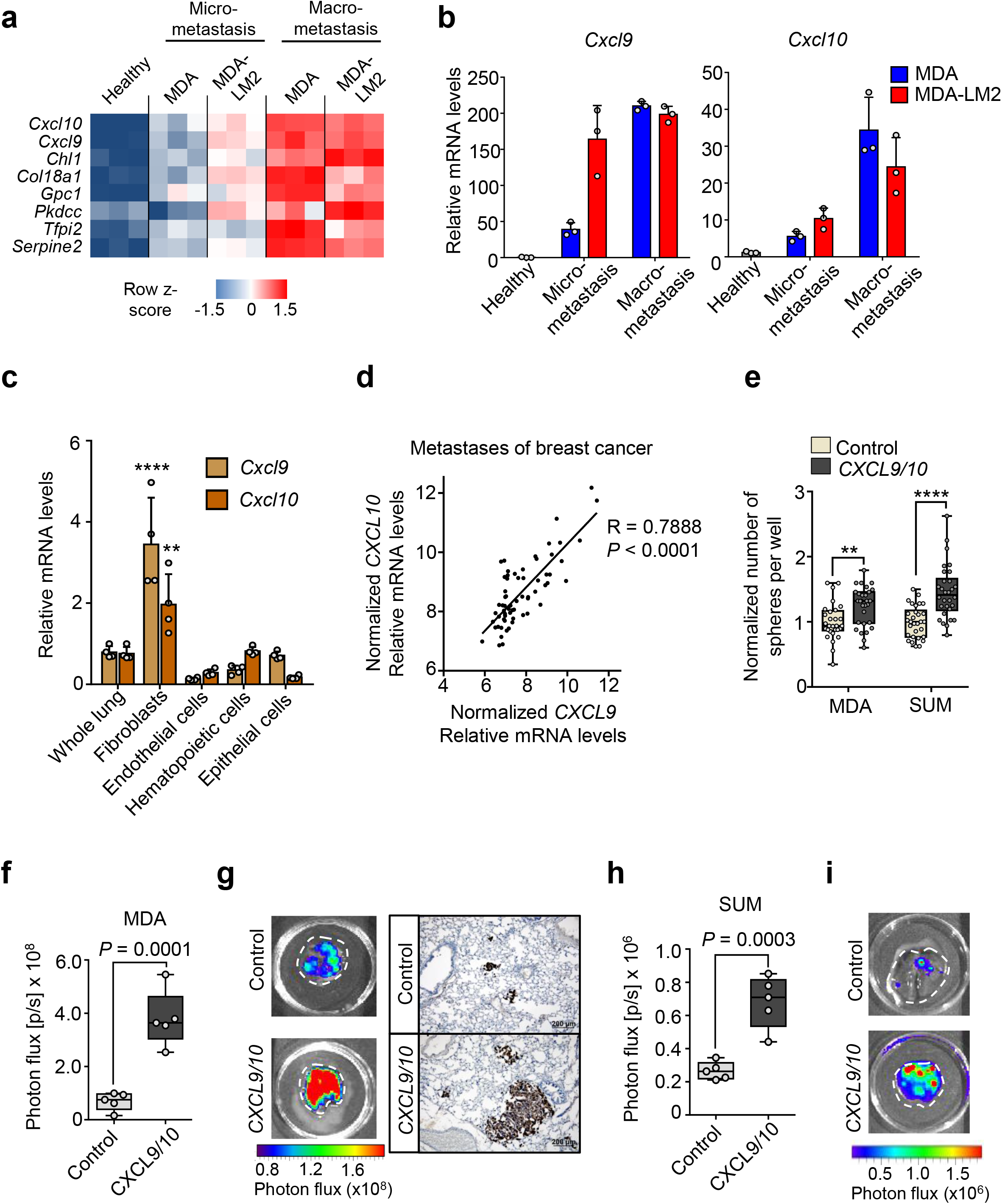
CXCL9/10 expression is induced in metastasis-associated fibroblasts and promotes metastatic colonization. **a**, Heatmap showing normalized mRNA expression of genes encoding secreted proteins that are induced in fibroblasts from MDA-LM2 micrometastasis and further induced in MDA- and MDA-LM2 macrometastasis. **b**, RT-qPCR validation of *Cxcl9* and *Cxcl10* expression in isolated fibroblasts from MDA- and MDA-LM2-induced lung metastases. Shown are means from three mice per group with standard deviation. **c**, *Cxcl9* and *Cxcl10* expression in indicated cell types isolated from MDA-LM2-macrometastatic lungs relative to overall expression in whole lungs. Gene expression was analyzed by RT-qPCR. *P* values were calculated by two-way ANOVA with Tukey’s multiple comparisons test to compare *Cxcl9/10* expression in all indicated cell types. Shown are summarized *P* values of *Cxcl9/10* upregulation in fibroblasts compared to all other cell types. Individual *P* values for expression of *Cxcl9/Cxcl10* in fibroblasts versus indicated cell populations are: versus whole lung: *P* < 0.0001 / *P* = 0.0048, versus endothelial cells: *P* < 0.0001 / *P* < 0.0001, versus hematopoietic cells: *P* < 0.0001 / *P* = 0.0080, versus hematopoietic cells: *P* < 0.0001 / *P* < 0.0080, versus epithelial cells: *P* < 0.0001 / *P* < 0.0001. *P* values in all other comparisons were not significant (*P* > 0.05). Data show means from four mice with standard deviation. **d**, Correlation analysis of *CXCL9* and *CXCL10* expression in dissected human metastases from breast cancer patients (GSE14020). Linear regression with Pearson correlation r and two-tailed *P* value, *n* = 65. **e**, Oncosphere formation of MDA or SUM breast cancer cells overexpressing *CXCL9/10* or a vector control. Data represent 3 independent experiments with quantification of 10 wells per condition. Boxes depict median with upper and lower quartiles. Whiskers show minimum to maximum and dots indicate each individual data point. *P* values were calculated by unpaired two-tailed t-tests. **f**, Lung colonization determined by bioluminescence in mice 16 days after intravenous injection of MDA breast cancer cells overexpressing *CXCL9/10* or a vector control. *P* value was calculated by an unpaired one-tailed t-test, *n* = 5 mice per group. **g**, Examples of *ex vivo* bioluminescence (left) and histology (right, Vimentin staining) from lungs of mice injected with MDA cells overexpressing *CXCL9/10* or a control vector. **h**, Lung colonization determined by bioluminescence in mice 16 days after intravenous injection of SUM breast cancer cells overexpressing *CXCL9/10* or a vector control. *P* value was calculated by an unpaired one-tailed t-test. **i**, Examples of *ex vivo* bioluminescence from lungs of mice injected with SUM cells overexpressing *CXCL9/10* or a control vector.

Among the earliest and most highly upregulated genes encoding secreted proteins were the two inflammatory chemokine (C-X-C motif) ligands 9 and 10 (*Cxcl9/10*) (Fig. 2a,b and Supplementary Fig. 5a,b). Expression analysis of different cell types isolated from lungs with growing metastases revealed that fibroblasts are the main source of *Cxcl9* and *Cxcl10* (Fig. 2c). Importantly, *CXCL9* and *CXCL10* expression strongly correlated in data sets of dissected distant metastases samples from breast cancer patients (Fig. 2d), indicating co-expression of the cytokines in human metastasis. Therefore, to address the functional role of CXCL9 and CXCL10, we ectopically expressed the genes together in parental MDA breast cancer cells as well as in SUM159 (SUM) cells, a second human breast cancer cell line (Supplementary Fig. 5c). Combined overexpression of *CXCL9* and *CXCL10* in MDA and SUM parental breast cancer cells significantly increased their ability to form spheres when cultured under serum-free low adhesive conditions compared to control cells (Fig. 2e). These results suggested that CXCL9 and CXCL10 can confer stem cell properties to breast cancer cells, as the ability to form oncospheres is associated with stem cell features in cultured cells^20^. To analyze the role of these chemokines in metastasis, we intravenously injected breast cancer cells co-expressing *CXCL9* and *CXCL10* into female, non-obese, diabetic-severe combined immunodeficiency gamma (NSG) mice. Notably, ectopic expression of *CXCL9* and *CXCL10* significantly promoted lung colonization by both MDA and SUM breast cancer cells (Fig. 2f-i). To address whether both *CXCL9* and *CXCL10* contribute to oncosphere formation and lung metastasis, we overexpressed the genes individually in MDA or SUM cancer cells (Supplementary Fig. 5d). Indeed, both *CXCL9* and *CXCL10* promoted sphere formation and lung colonization by breast cancer cells (Supplementary Fig. 5e,f). Together, these data show that *CXCL9* and *CXCL10* represent components of the metastatic niche that are co-induced in activated MAFs in lungs and support lung colonization.

### Metastatic breast cancer cells secrete IL-1α and IL-1β that induce *CXCL9* and *CXCL10* expression in stromal lung fibroblasts by an NF-κB-dependent mechanism

We next examined how *Cxcl9 and CXCL10* are induced in lung fibroblasts during metastatic colonization and whether breast cancer cells directly account for this induction. Gene set enrichment analysis (GSEA) revealed a significant enrichment of pro-inflammatory Interleukin-1 (IL-1) cytokine response and NF-κB signaling in fibroblasts from MDA- and MDA-LM2-derived macrometastases (Fig. 1f and 3a; and Supplementary Table 1). Notably, an enrichment of this signaling was observed already at a micrometastatic stage in MDA-LM2-associated fibroblasts, thus correlating with the observed induction of *Cxcl9/10* (Fig. 1f, 2a and 3a; and Supplementary Table 1). In line with this, we observed a significant correlation between *CXCL9/10* and *IL1A/B* expression in dissected metastases samples from breast cancer patients (Fig. 3b). Therefore, we hypothesized that IL-1α/β present in metastatic lungs may induce *Cxcl9/10* expression in lung fibroblasts. Indeed, stimulation with recombinant IL-1α and IL-1β significantly induced expression of *CXCL9* and *CXCL10* in MRC-5 human lung fibroblasts, and this induction was mediated by NF-κB activity (Fig. 3c and 3d). Moreover, blockade of IL-1 receptor (IL-1R) signaling through the use of an inhibitory human anti-IL-1R monoclonal antibody blunted the induction of *CXCL9* and *CXCL10* in fibroblasts by recombinant IL-1α and IL-1β (Supplementary Fig. 6a,b). These results indicated that IL-1α and IL-1β mediated induction of CXCL9 and CXCL10 in the fibroblasts via activation of IL-1 receptor and downstream NF-κB activation.

**Fig. 3.**
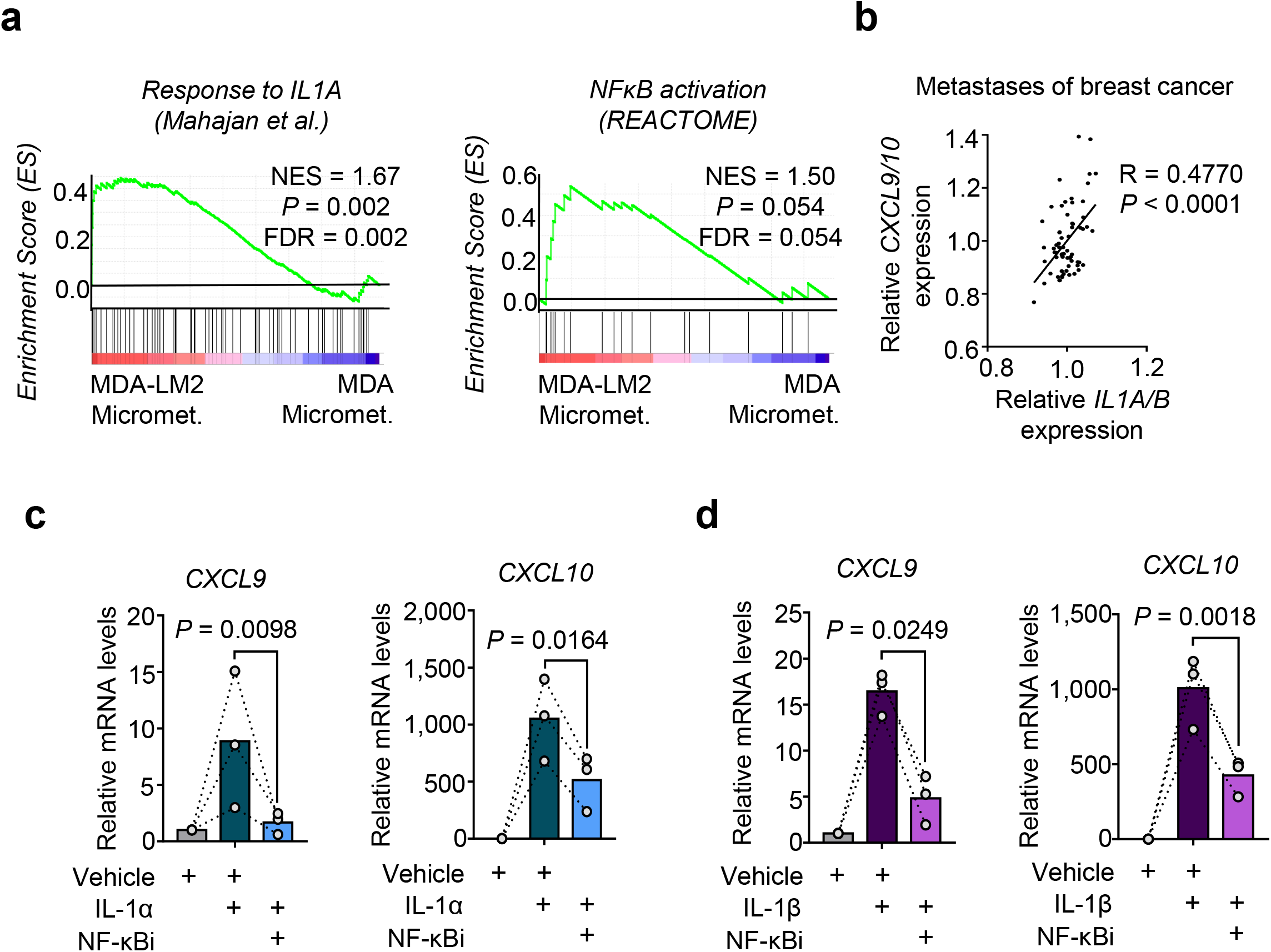
*CXCL9/10* expression in fibroblasts is induced by IL-1α/β via NF-κB signaling. **a**, Enrichment of IL-1α and NF-κB signatures in fibroblasts isolated from lungs bearing MDA-LM2 micrometastases compared to MDA parental micrometastases. NES, normalized enrichment score, FDR, false discovery rate. *P* values were determined by random permutation tests. **b**, Correlation analysis of mean *CXCL9/10* and mean *IL1A/B* expression in data sets from human breast cancer metastases. Linear regression with Pearson correlation r and two-tailed *P* value. *n* = 65. **c-d**, *CXCL9* and *CXCL10* expression in MRC-5 human lung fibroblasts treated with 1 ng/ml recombinant human IL-1α (c) or IL-1β (d) alone or in combination with 5 μM JSH-23 (NF-κBi) for 48 h. Expression was analyzed by RT-qPCR. *P* values were determined by ratio-paired one-tailed t-tests; *n* = 3 independent experiments.

We hypothesized that the cancer cells may be a direct source of the IL-1 ligands that induce *CXCL9/10* in MAFs. Indeed, *IL1A* and *IL1B* were expressed by MDA and SUM breast cancer cell lines, and *IL1A/B* expression levels were significantly increased in the respective lung metastatic derivatives MDA-LM2 and SUM-LM1, suggesting an association with metastatic potential (Fig. 4a). To measure protein levels of cancer cell-derived IL-1α and IL-1β *in situ*, we carried out human-specific enzyme-linked immunosorbent assays (ELISAs) on whole lung homogenates from mice bearing MDA-LM2-derived lung metastases. ELISAs confirmed expression of both cytokines in metastatic lungs (Fig. 4b). Considering IL-1α and IL-1β are secreted cytokines, we investigated whether conditioned medium (CM) of cultured breast cancer cells can drive induction of *CXCL9/10* in fibroblasts *in vitro*. We treated MRC-5 cells with CM from parental (MDA/SUM) or highly metastatic (MDA-LM2/SUM-LM1) breast cancer cells (Fig. 4c). In line with the observed high levels of *IL1A/B* in metastatic breast cancer cells (Fig. 4a), treatment with CM from MDA-LM2 or SUM-LM1 cancer cells induced a stronger upregulation of *CXCL10* in MRC-5 fibroblasts than CM from MDA or SUM parental cell counterparts (Fig. 4d). Moreover, addition of a blocking antibody against IL-1R1 or an NF-κB inhibitor to the CM of MDA-LM2 and SUM-LM1 cells prevented induction of *CXCL10* in MRC-5 fibroblasts (Fig. 4e,f and Supplementary Fig. 6c). Importantly, treatment of lung fibroblasts with CM from MDA-LM2 cells transduced with shRNA against IL-1α/β or treatment of IL-1R knockout (IL1R-KO) fibroblasts with CM from MDA-LM2 cells also prevented upregulation of *CXCL10* in the fibroblasts (Fig. 4g-i, Supplementary Fig. 6d). These experiments confirm that IL-1α/β are indeed the factors contained within the CM from metastatic breast cancer cells that drive *CXCL10* expression in lung fibroblasts. Importantly, similar effects were observed when we stimulated fibroblasts with CM from patient-derived cancer cells that were collected from pleural effusions of advanced breast cancer patients, as this CM induced *CXCL10* in MRC-5 cells in an NF-κB-dependent manner (Fig. 4j,k). To address whether IL-1 cytokines play a functional role in metastatic colonization of the lungs, we injected control and shIL1A/B transduced MDA-LM2 cells intravenously into NSG mice and measured lung colonization by bioluminescence. IL-1α/β knockdown cancer cells showed a significantly reduced ability to colonize the lung, indicating that the IL-1 cytokines are required for the growth of lung metastasis (Fig. 4l,m). Together, these findings suggest that IL-1α/β secreted by breast cancer cells that have reached the lungs directly activate IL-1R on lung fibroblasts to induce NF-κB-dependent *CXCL9/10* expression that promotes lung colonization.

**Fig. 4.**
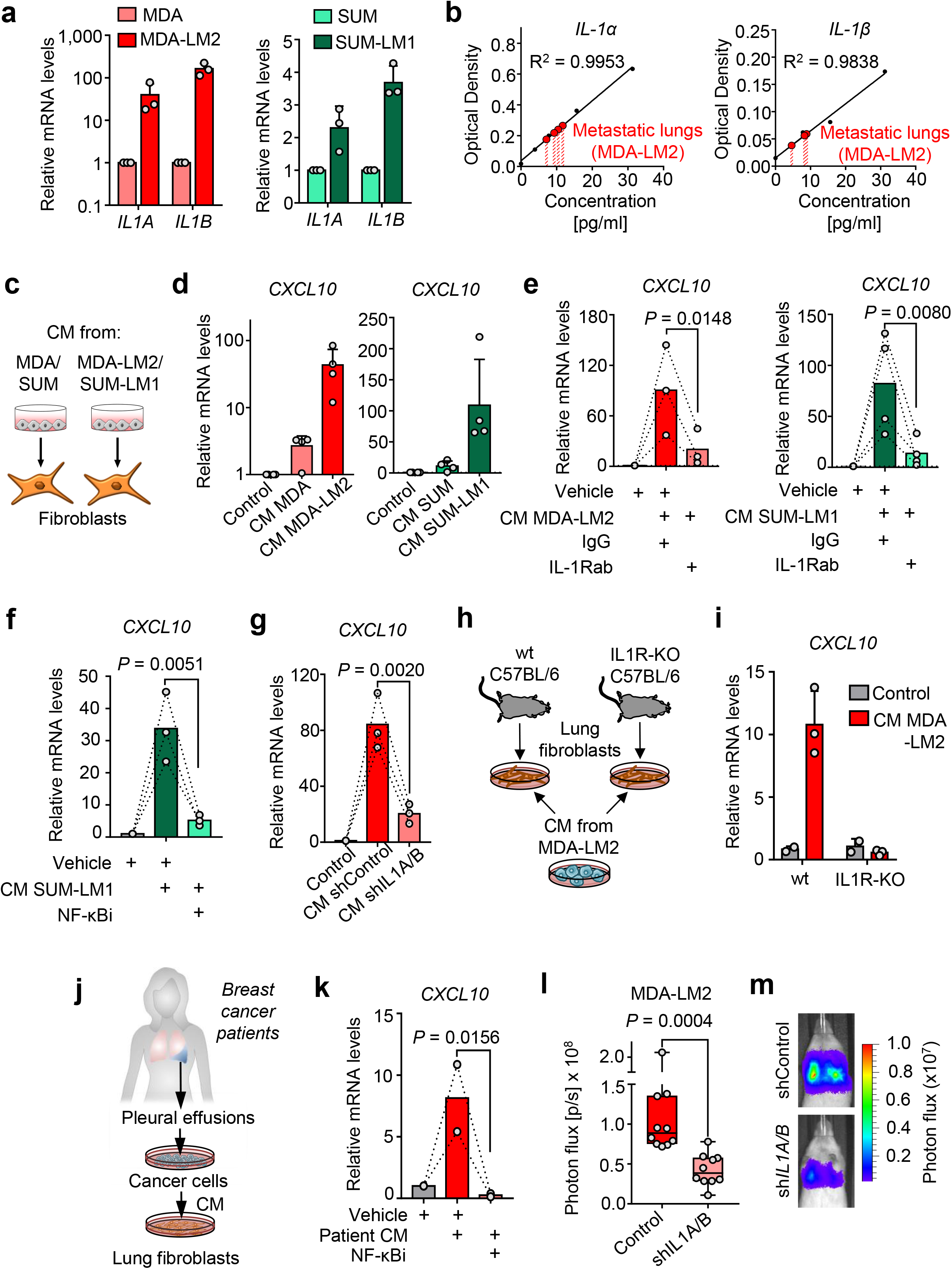
Metastatic breast cancer cells express IL-1α and IL-1β to induce *CXCL9/10* in reactive fibroblasts. **a**, *IL1A* and *IL1B* expression in MDA and SUM parental breast cancer cells and their metastatic derivatives MDA-LM2 and SUM-LM1. Expression was analyzed by RT-qPCR; *n* = 3. **b**, Human IL-1α and IL-1β protein levels in metastatic lungs from xenograft mouse models after intravenous injection of MDA-LM2 cancer cells. Cytokine levels were measured by ELISA. Shown are standard curves and interpolated IL-1α and IL-1β concentrations in lung metastasis lysates from 4 mice (IL-1α) and 3 mice (IL-1β). Samples were dilute 1:2, thus final concentrations per lung are twice as high. **c**, Schematic of setup for experiments using cancer cell conditioned medium to stimulate fibroblasts. **d**, *CXCL10* expression in MRC-5 fibroblasts treated with conditioned medium (CM) from MDA/SUM parental breast cancer cells or their metastatic derivatives MDA-LM2/SUM-LM1 for 48 h, *n* = 4 experiments. **e-f**, *CXCL10* expression in MRC-5 fibroblasts in response to MDA-LM2 or SUM-LM1 cancer cell CM alone or co-treated with 20 μg/ml IL-1R1 blocking antibody or isotype control (e) or 5 μM JSH-23 (NF-κBi) or vehicle (f) for 48 h. MDA-LM2, *n* = 3; SUM-LM1, *n* = 4; vehicle, *n* = 3. **g**, *CXCL10* expression in fibroblasts treated with CM from control or *IL1A/B* knockdown MDA-LM2 cancer cells for 48h, *n* = 3 experiments. **h-i**, Fibroblasts isolated from lungs of wt and IL-1R KO C57BL/6 mice and treated with MDA-LM2 conditioned medium (h). *CXCL10* mRNA levels in CM-treated wt and IL-1 KO fibroblasts (i). Expression was determined by RT-qPCR. **j-k**, *CXCL10* expression in fibroblasts treated with CM from primary cancer cells derived from pleural effusions of metastatic breast cancer patients alone or in combination with 5 μM JSH-23 (NF-κBi) for 48 h. *P* values in panels (e-h) were determined by ratio-paired one-tailed t-tests. **l**, Lung colonization in mice injected intravenously with control or shIL1A/B transduced MDA-LM2 cancer cells as determined by bioluminescence. *P* values were calculated by unpaired one-tailed t-tests; *n* = 10 mice per group. **m**, Representative luminescence images from each group in panel (l).

### Expression of *IL1A/B* in metastatic breast cancer cells is driven by JNK activity

We previously demonstrated that the JNK signaling pathway promotes lung metastasis via induction of the ECM proteins osteopontin (SPP1) and tenascin C (TNC)^21^. These studies also revealed that JNK activity in breast cancer cells promotes the expression of *IL1A* and *IL1B*. As expression levels of both *IL1A* and *IL1B* were significantly higher in MDA-LM2 cells compared to their parental line (Fig. 4a), we hypothesized that this may be explained by a higher JNK activity in the metastatic derivative MDA-LM2. Indeed, the JNK response signature^21^ was significantly enriched in highly metastatic MDA-LM2 cells compared to MDA parental cells, both *in vivo* and *in vitro* (Fig. 5a and Supplementary Fig. 6e). To confirm regulation of *IL1A* and *IL1B* by JNK in breast cancer cells, we determined their mRNA levels in MDA-LM2 cells upon expression of a constitutively active form of JNK, consisting of a protein fusion between JNK1 and its upstream MAPK kinase (MAPKK) activator MKK7 (MKK7-JNK), or a mutated version (MKK7-JNK(mut)), in which the phosphorylation motif Thr180-Pro-Tyr182 in JNK1 is replaced with Ala-Pro-Phe, thereby preventing its activation by MKK7^22, 23^. In line with our previous observations ^21^, *MKK7-JNK* expression significantly induced both *IL1A* and *IL1B*, and this induction was blunted in *MKK7-JNK(mut)-expressing* cells (Fig. 5b). Moreover, treatment with a JNK inhibitor (JNKi) significantly reduced endogenous *IL1A* and *IL1B* expression in MDA-LM2 cells (Fig. 5c). To determine whether JNK induces *IL1A* and *IL1B* via the transcription factor c-Jun, we performed chromatin immunoprecipitation (ChIP), pulling down c-Jun and bound chromatin. qPCR analysis of c-Jun-bound chromatin confirmed that c-Jun binds to *IL1A* and *IL1B* promoters in breast cancer cells (Fig. 5d,e and Supplementary Fig. 6f).

**Fig. 5.**
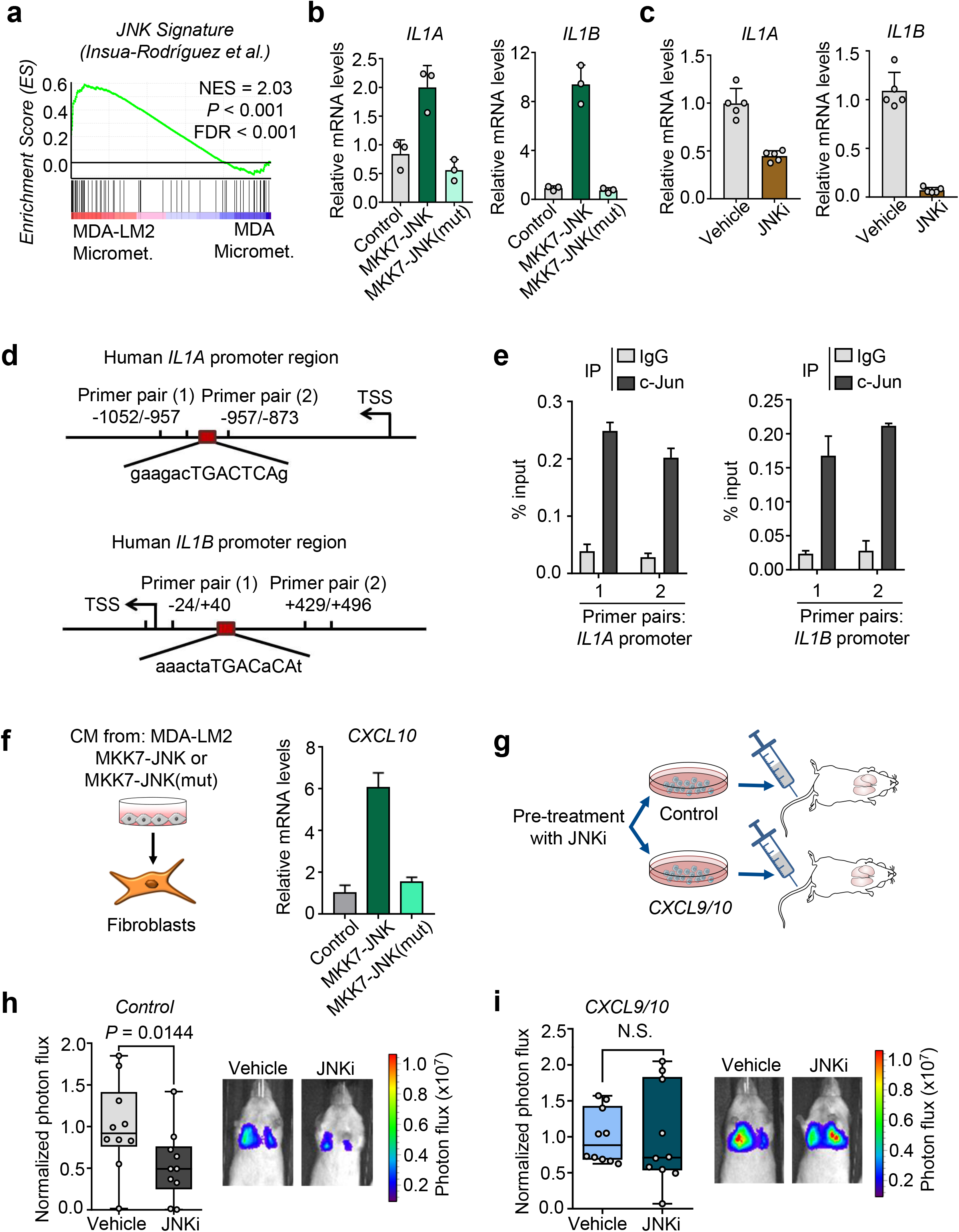
JNK signaling in breast cancer cells induces *IL1A/B*. **a**, Enrichment of a JNK response signature ^21^ in MDA-LM2 cells compared to parental MDA cells isolated from lung micrometastases (transcriptome from cells *in situ*). NES, normalized enrichment score, FDR, false discovery rate. *P* value was determined by random permutation test. **b**, Expression of *IL1A* and *IL1B* in MDA-LM2 breast cancer cells expressing activated JNK (MKK7-JNK) or a mutated version of JNK (MKK7-JNK(mut)) or vector control (Control) determined by qPCR, *n* = 3 experiments. **c**, *IL1A/B* expression in MDA-LM2 cells treated with 5 μM CC-401 JNK inhibitor (JNKi) or vehicle, *n* = 5 experiments. **d**, Maps of *IL1A* and *IL1B* promoter regions showing c-Jun consensus binding sites and primer positions used to perform CHIP-qPCR. **e**, CHIP analysis where c-Jun bound to chromatin was pulled down and *IL1A* and *IL1B* promoter chromatin was analyzed by qPCR in MDA-LM2 cells. **f**, *CXCL10* expression in MRC-5 fibroblasts treated with conditioned medium from MDA-LM2 breast cancer cells expressing MKK7-JNK fusion gene, MKK7-JNK(mut) or a control vector (Control). Gene expression in (b,c and f) was determined by RT-qPCR. Results are representative of two independent experiments. **g**, Diagram of metastasis experiments where MDA-LM2 breast cancer cells, expressing ectopic *CXCL9/10* (*CXCL9/10*) or a control vector (Control) were treated with 5 μM CC-401 (JNKi) or vehicle for 48 h before intravenous injection into NSG mice for lung colonization. **h-i**, Lung colonization of Control (h) or *CXCL9/10* overexpressing MDA-LM2 cancer cells (i) pretreated with 5 μM JNKi or vehicle for 48 h. Lung colonization was determined by bioluminescence. Shown are photon flux quantification and representative bioluminescence images. *P* values were calculated by unpaired one-tailed t-tests; *n* = 10 mice per group.

Consistent with JNK-driven expression of *IL1A/B* in cancer cells, CM from *MKK7-JNK*-expressing MDA-LM2 breast cancer cells increased the production of *CXCL10* in fibroblasts compared to CM control, and this increase was blunted when fibroblasts were treated with CM from *MKK7-JNK(mut)*-expressing cells (Fig. 5f). Importantly, these findings suggested that inhibition of JNK activity in breast cancer cells may alter their ability to induce a pro-metastatic paracrine crosstalk with lung fibroblasts. To test this hypothesis, we pre-treated MDA-LM2 cells overexpressing *CXCL9* and *CXCL10* or a control vector with JNKi and injected the cells intravenously into NSG mice (Fig. 5g and Supplementary Fig. 6g). In mice injected with MDA-LM2 control cells, pre-treatment with JNKi significantly reduced metastatic colonization (Fig. 5h and Ref.^21^). However, in mice injected with cancer cells overexpressing *CXCL9* and *CXCL10*, no reduction in metastasis was observed in response to JNKi treatment (Fig. 5i). These results indicate that JNK-driven production of IL-1α/β by metastatic cancer cells induces CXCL9/10 in pulmonary MAFs to form a supportive metastatic niche.

### CXCR3 marks a subset of breast cancer cells that can both induce and benefit from the pro-metastatic crosstalk with lung fibroblasts

The G-protein coupled receptor CXCR3 is the only cellular receptor known to bind and become activated by CXCL9 and CXCL10^24^. Interestingly, flow cytometric analysis revealed that CXCR3 is expressed in a subpopulation of MDA and SUM cancer cells as well as their metastatic derivatives (Fig. 6a). Moreover, the proportion of CXCR3^+^ cancer cells was significantly higher when cultured in serum-free sphere conditions, and was higher in the lung metastatic derivative SUM-LM1 compared to the respective parental counterpart (Fig. 6a). Importantly, CXCR3 was also expressed in subsets (range 3.8% - 11.1%) of cancer cells isolated from pleural effusions or ascites of four breast cancer patients (Supplementary Fig. 7a). To further characterize the CXCR3-expressing subpopulation of breast cancer cells, we established transcriptomic profiles of FACS-sorted CXCR3^+^ and CXCR3^−^ SUM-LM1 breast cancer cells (Fig. 6b and Supplementary Table 2). Intriguingly, GSEA revealed that CXCR3^+^ cancer cells had increased inflammatory signaling and higher JNK activity, and showed characteristics of basal and stem cells of the mammary gland (Fig. 6c,d and Supplementary Fig. 7b,c), in line with the observed increase of CXCR3^+^ cells in sphere cultures (Fig. 6a). GO term analysis further indicated an enrichment of genes involved in inflammatory signaling and chemokine production (Supplementary Table 3). We therefore reasoned that CXCR3^+^ cancer cells may secrete higher levels of IL-1α/β, which could in turn lead to elevated production of *CXCL9* and *CXCL10* in fibroblasts. Indeed, isolated CXCR3^+^ cancer cells expressed higher levels of *IL1A* and *IL1B* compared to CXCR3^−^ counter parts (Fig. 6e,f), and CM from CXCR3^+^ 4T1 mouse mammary tumor cells, but not from CXCR3^−^ cells, induced *CXCL10* expression in mouse and human fibroblasts (Fig. 6g and Supplementary Fig. 7d). Importantly, sorted CXCR3^+^ 4T1 mammary cancer cells had a higher tumor-initiating ability compared to sorted CXCR3^−^ cells when co-injected with lung fibroblasts, subcutaneously in limiting dilutions, and resulting tumors by CXCR3^+^ cells were significantly larger (Fig. 6h–j). Furthermore, sorted CXCR3^+^ MDA-LM2 cells had significantly increased abilities to establish metastases in the lung microenvironment compared to CXCR3^−^ cells (Fig. 6k,l). Collectively, these data indicate that CXCR3^+^ metastasis-initiating cells can induce a fibroblast niche in the lung and underscore the importance of a paracrine interaction with fibroblasts in promoting tumor and metastasis initiation.

**Fig. 6.**
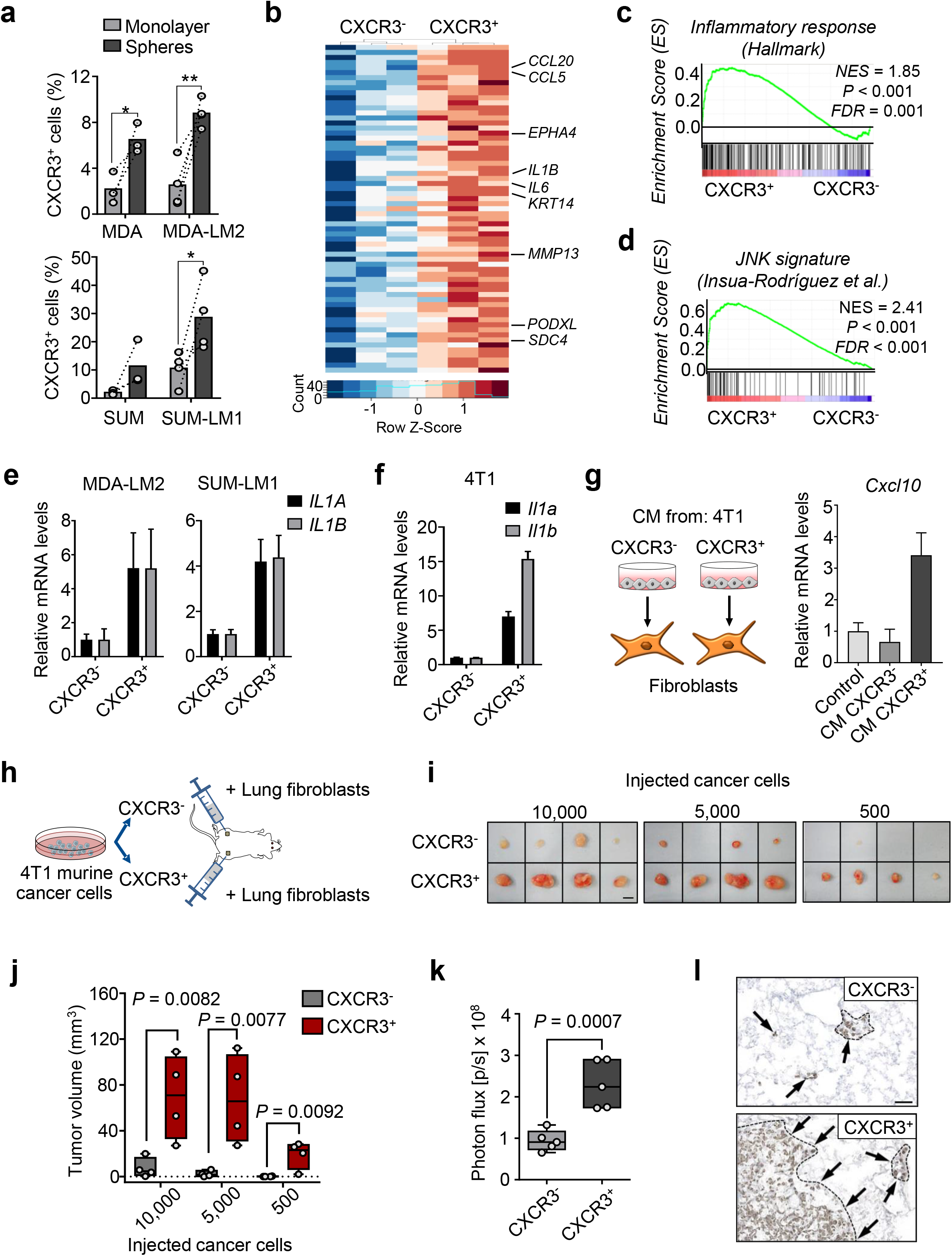
CXCR3^+^ breast cancer cells induce and benefit from paracrine crosstalk with lung fibroblasts. **a**, Proportion of CXCR3-expressing MDA, MDA-LM2, SUM and SUM-LM1 breast cancer cells in monolayer or oncosphere cultures as determined by flow cytometric staining. *P* values were calculated by paired one-tailed t-tests. MDA and SUM, *n* = 3 independent experiments; MDA-LM2 and SUM-LM1, *n* = 4 independent experiments. **b**, Heatmap showing normalized expression of genes induced in sorted CXCR3-positive (CXCR3^+^) compared to CXCR3-negative (CXCR3^−^) SUM-LM1 breast cancer cells. Shown are selected genes induced in CXCR3^+^ population. **c-d**, Enrichment of indicated gene sets in CXCR3^+^ SUM-LM1 breast cancer cells compared to CXCR3^−^ cells. NES, normalized enrichment score, FDR, false discovery rate. *P* values were determined by random permutation tests. **e-f**, *IL1A/B* expression in sorted CXCR3^+^ human breast cancer cells (MDA-LM2 and SUM-LM1) (e) and *Il1a/b* expression in mouse mammary tumors cells (4T1) (f), respectively. RT-qPCR data is representative of 3 independent replicates each. **g**, *Cxcl10* expression in NSG mouse lung fibroblasts treated with control medium or conditioned medium (CM) from CXCR3^+^ and CXCR3^−^ 4T1 mammary tumor cells for 48 h. **h**, Diagram of tumor initiation experiment in mice where sorted CXCR3^+^ or CXCR3^−^ 4T1 tumor cells were co-injected subcutaneously with primary mouse lung fibroblasts into BALB/c mice in limiting dilutions on either flank of the mouse. **i**, Subcutaneous tumors resected 3 weeks after injection of CXCR3^+^ and CXCR3^−^ 4T1 breast cancer cells and lung fibroblasts injected in limiting dilution into mice as described in panel (h); *n* = 4 mice per group. Scale bar 5 mm. **j**, Quantification of tumor sizes from the experiment described in panels (h,j). *P* values were calculated by unpaired one-tailed t-tests. **k**, Bioluminescence analysis of lung colonization of sorted CXCR3^+^ or CXCR3^−^ MDA-LM2 breast cancer cells 16 days after intravenous injection into NSG mice. *P* value was calculated by an unpaired one-tailed t-test; *n* = 5 mice per group. **l**, Histological examples of metastases determined by expression of human vimentin in lungs of mice injected with CXCR3^+^ or CXCR3^−^ MDA-LM2 breast cancer cells. Scale bar, 50 μM.

### Inhibition of CXCR3 blocks lung colonization in xenograft and syngeneic mouse models

In addition to the increased ability of CXCR3^+^ cancer cells to induce *Cxcl9/10* expression in lung fibroblasts, this subpopulation of cancer cells is also likely to benefit from this crosstalk. To test whether CXCR3 is functionally required for the *CXCL9/10*-mediated increase in metastatic ability in breast cancer cells, we used the CXCR3 antagonist AMG-487 (CXCR3i). Stimulation of MDA-LM2 cells, SUM159-LM1 cells and patient-derived breast cancer cells with recombinant *CXCL9* and/or *CXCL10* increased sphere formation, and this was reversed by addition of CXCR3i (Fig. 7a,b). Importantly, systemic treatment of NSG mice with CXCR3i significantly diminished lung metastatic outgrowth of MDA-LM2 cells (Fig. 7c). Since CXCL9 and CXCL10 are known to function in the regulation of immune responses^25, 26^, we tested the effect of CXCR3i in an immunocompetent mouse model. BALB/c mice were injected intravenously with 4T1 mouse mammary tumor cells and concurrently treated with CXCR3i until the experimental endpoint. Lung metastatic outgrowth was also significantly reduced upon CXCR3i in the syngeneic setting (Fig. 7d,e), indicating that systemic antagonism of CXCR3 may be an effective strategy to disrupt cancer cell-fibroblast crosstalk that fuels metastatic colonization of the lungs. To address the putative link between CXCR3-expressing cancer cells and clinical prognosis, we clustered samples from breast cancer patients according to the expression of CXCR3 signature (CXCR3S) (Supplementary Table 2). Analysis of the TOP trial data set and a collection of independent data sets from basal-like breast cancers, revealed that patients with high CXCR3S associated with poor relapse-free survival, distant metastasis-free survival and overall survival (Fig. 7f,g; Supplementary Fig. 8). Taken together, these evidences suggest that CXCR3^+^ metastasis-initiating cells not only induce CXCL9 and CXCL10 in MAFs, but also take advantage of these cytokines to promote metastatic colonization.

**Fig. 7.**
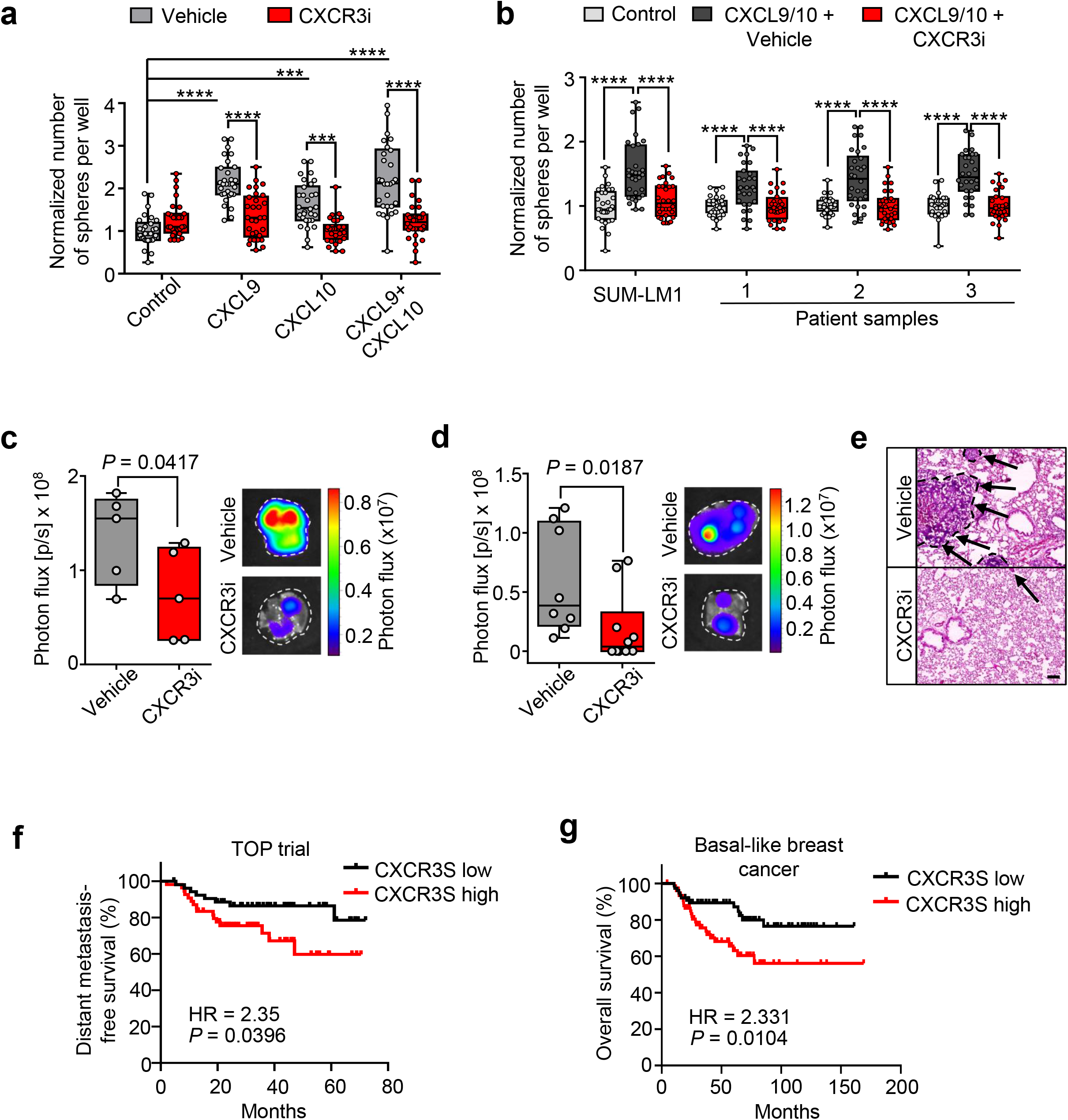
CXCR3 mediates CXCL9/10-induced oncosphere formation and can be targeted to inhibit lung metastasis. **a**, Quantification of oncospheres per well from MDA-LM2 cells stimulated with 100 ng/ml recombinant CXCL9 or CXCL10 individually or together in combination with 10 μM CXCR3 antagonist AMG-487 (CXCR3i) or vehicle control (Vehicle) for 7 days. **b**, Quantification of oncospheres derived from SUM-LM1 breast cancer cells and primary breast cancer cells from patient samples isolated from pleural fluids or ascites upon stimulation with 100 ng/ml recombinant CXCL9/10 in combination with CXCR3i or Vehicle for 7 days. For (a) and (b), sphere numbers per well were normalized to the average number in the control group. Data represent 3 independent experiments with quantification of 9-10 wells per condition. Boxes depict median with upper and lower quartiles. Whiskers show minimum to maximum and dots indicate each individual data point. *P* values were calculated by ordinary one-way ANOVA with Tukey’s multiple comparisons test. **c-d**, Lung colonization upon systemic treatment with CXCR3i in xenograft and syngeneic mouse models. MDA-LM2 breast cancer cells (c) or 4T1 mouse mammary tumor cells (d) were injected intravenously into NSG and BALB/c mice, respectively, and mice received 8 mg/kg AMG-487 (CXCR3i) twice daily for the duration of the experiment. Lung colonization was determined by bioluminescence 13 days after injection of cancer cells. Shown are photon flux quantification in lungs and representative *ex vivo* lung bioluminescence images. *P* values were calculated by unpaired one-tailed t-tests. MDA-LM2 analysis (c), *n* = 5 mice per group; 4T1 analysis (d) vehicle, *n* = 8 mice and CXCR3i, *n* = 10 mice (pooled from 2 independent experiment). **e**, Examples of metastases in H&E-stained lung from BALB/c mice after intravenous injection of 4T1 mammary cancer cells and continued treatment with CXCR3i or Vehicle. Scale bar, 100μm. **f-g**, Kaplan-Meier analyses of breast cancer patients, associating CXCR3^+^ cell signature (CXCR3S, mean expression of 65 genes) with distant metastasis-free survival (f, TOP trial data set, *n* = 107 patients) or overall survival (g, compiled data set from basal-like breast cancers, KM plotter, *n* = 153). Median cutoff was used to group samples into CXCR3S low and high. HR, hazard ratio. *P* values were determined by log-rank test.

## DISCUSSION

Disseminated cancer cells require a supportive niche to successfully form metastases^5^. Indeed, the vast majority of cancer cells face an unfavorable microenvironment at secondary sites and are eliminated following extravasation^27, 28^. Resting stroma can be highly resistant to the establishment of intruding cells^29^, so cancer cells that arrive at distant organs equipped with their own niche components or niche-promoting ability are likely to have a selective advantage. Our earlier work provides an example of how cancer cells can benefit from bringing own niche components^15^. In this study, we show how enhanced niche-promoting ability fuels metastatic colonization.

JNK signaling promotes metastasis of breast cancer cells through distinct mechanisms. Our previous study showed that successful breast cancer metastasis requires JNK-induced expression of the extracellular matrix and niche proteins SPP1 and TNC^21^. Here, we reveal that JNK signaling also promotes communication between cancer cells and lung fibroblasts and enables highly metastatic breast cancer cells to rapidly establish a supportive niche in the lung. Our findings suggest a model (summarized in Fig. 8), in which high JNK activity in metastasis-initiating breast cancer cells induces the expression of IL-1α and IL-1β that interact with IL-1R on stromal fibroblasts in the lung to stimulate an NF-κB-mediated induction of *Cxcl9* and *Cxcl10*. Once secreted from the fibroblasts, CXCL9 and CXCL10 bind to CXCR3 receptor on the surface of a subpopulation of breast cancer cells to complete a paracrine loop between the two cell types that promotes growth of breast cancer metastases. Our results establish a link between stress signaling, the ability of disseminated cancer cells to modify the microenvironment in secondary organs, and their metastatic potential.

**Fig. 8.**
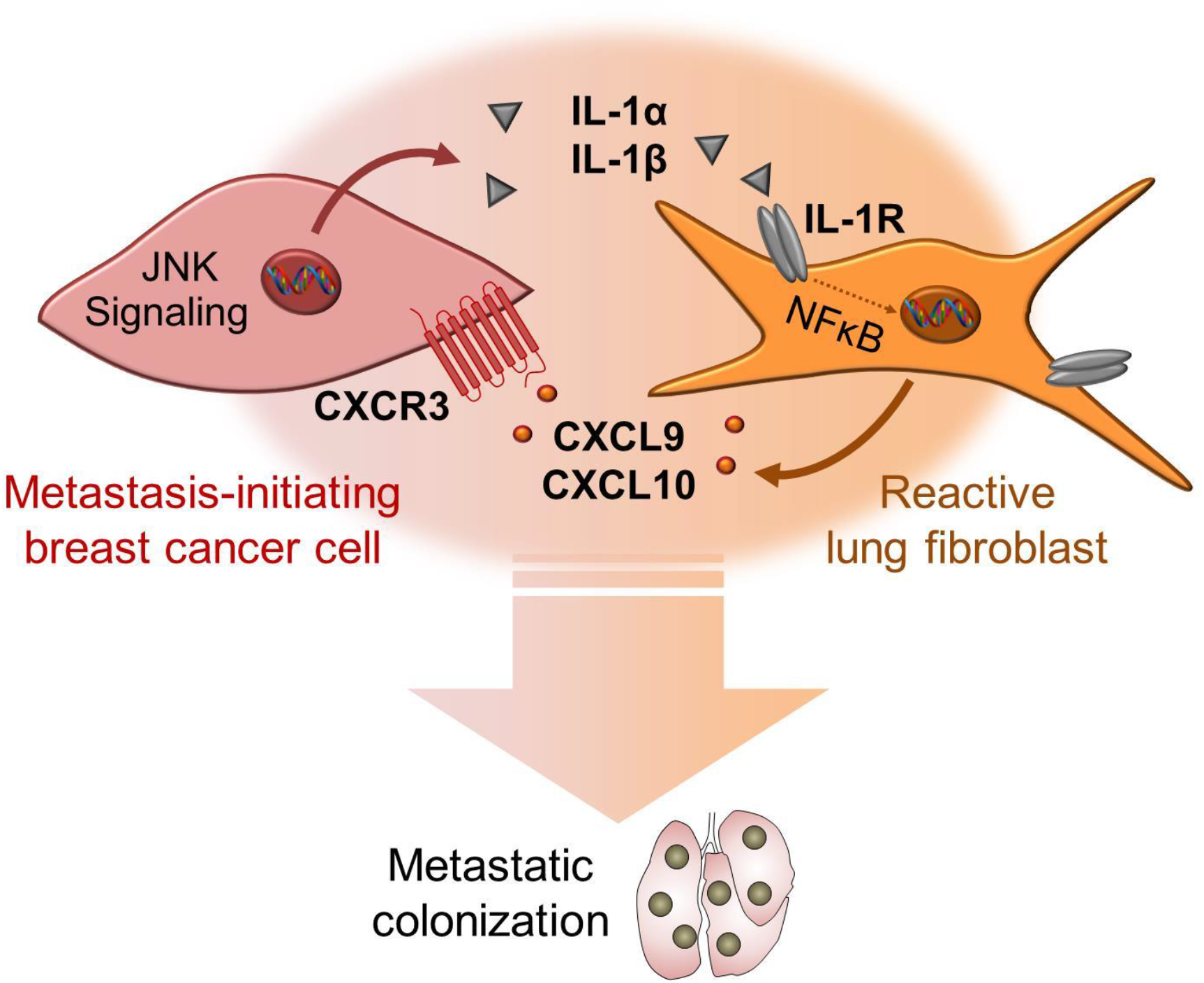
Model of interaction between metastasis-initiating breast cancer cells and fibroblasts in the lungs. CXCR3^+^ breast cancer cells exhibit high JNK activity that drives secretion of IL-1α/β. Fibroblasts respond to IL-1α/β via type I IL-1R signaling and NF-kB-mediated upregulation of CXCL9/10 that promote metastatic colonization via CXCR3.

The role of inflammatory signaling in cancer is complex and is likely to be context-dependent. At primary sites, IL-1 signaling has been shown to promote tumor growth^10, 30^. However, in secondary organs studies have suggested both a pro-metastatic and anti-metastatic roles for IL-1 signaling in models of breast cancer^31, 32^. The divergent IL-1 responses in metastasis may be explained by different breast cancer subtypes in focus. Our study is focused on basal-like breast cancer that has high propensity to metastasize to lung. Inflammatory signaling such as NF-κB is active in basal-like breast cancer^33, 34^, indicating that this breast cancer subtype can adapt to and take advantage of inflammatory signaling. Moreover, we have previously shown that JNK signaling, that induces IL-1α, IL-1β and other pro-metastatic factors in breast cancer cells, is particularly associated with basal-like breast cancer^21^. Finally, our results based on loss-of-function indicate that indeed IL-1, secreted by basal-like cancer cells, is required for metastatic colonization of the lung.

We find that reactive fibroblasts residing near or within metastatic lesions in the lung acquire an inflammatory phenotype reminiscent of the response of fibroblasts to wound healing and primary tumors^10, 35^. For example, collagen expression (particularly fibrillary collagens), extracellular matrix proteins (including fibronectin, TNC, and SPP1), and matrix-modifying enzymes (including serpins and lox-family proteins) are highly induced in MAFs. We find that this inflammatory phenotype is associated with a substantial expansion of fibroblasts, resulting in an approximate 50-fold increase in the number of fibroblasts in macrometastatic nodules compared to micrometastases, which is likely derived from the striking increase in proliferation that we observed within fibroblast populations. However, cancer- and metastasis-associated fibroblasts generally represent a heterogeneous group of mesenchymal cells, including resident tissue fibroblasts, pericytes and bone marrow-derived mesenchymal stromal cells^36^, and therefore this expanded population could include several subtypes of fibroblasts with unique functions and diverse origins. Recent studies suggest that indeed tumors may harbor a number of different fibroblast subtypes^37–39^. Certainly, fibroblasts go through distinct phases that are associated with inflammatory and contractile phenotypes during wound healing, but whether this is caused by changes to existing fibroblasts or by an influx of new fibroblast populations with different phenotypes is not well understood. Additional studies are therefore needed to determine whether the evolution of fibroblasts reacting to the metastatic stroma is mechanistically analogous to the changes that occur in fibroblasts during wound healing and whether it involves distinct subtypes.

Increasing evidence suggests that CXCR3 may play an important role during breast cancer progression^40–42^. We show that the ability to promote initiation of tumors and metastases is significantly enriched in CXCR3^+^ cancer cells compared to CXCR3^−^ cells. CXCR3^+^ breast cancer cells secrete IL-1α and IL-1β to stimulate a paracrine crosstalk with lung fibroblasts from which they benefit through their CXCR3 receptor. We find that only a small subset of metastatic breast cancer cells expresses CXCR3, and this subset is characterized by high JNK activity, which we previously linked to mammary stem cell properties^21^. Thus, CXCR3-expressing cancer cells may be enriched in metastatic stem cells that are equipped to exploit the metastasis-promoting paracrine loop between fibroblasts and cancer cells. This conclusion is also supported by previous work showing that cancer stem cells are able to take advantage of microenvironmental cues, such as the extracellular matrix protein periostin expressed by reactive lung fibroblasts, which supports stem cell maintenance and metastatic colonization^43^. Evidence from studies on colon cancer also indicates that fibroblasts play a major role in the maintenance of cancer stem cells^44, 45^. As an important addition to these reports, our findings indicate that metastatic stem cells may not only selectively exploit stromal signals at the distant site, but that they may also be selectively efficient in inducing the required stromal signals. Thus, CXCR3 expression may mark a unique population of breast cancer cells that strategically communicate with stromal fibroblasts to establish a supportive metastatic niche tailored to their phenotype/need.

Previous studies suggest a conflicting function for the CXCL9/10-CXCR3 axis during cancer progression that includes modulation of the immune microenvironment. Studies have shown that CXCL9/10-CXCR3 can mediate the recruitment and activation of T lymphocytes^25^ and in melanoma mouse models, this may lead to inhibition of tumor growth and progression^26^. However, in breast cancer, elevated CXCL10 levels are associated with increased recruitment of CXCR3-expressing, regulatory T cells and reduced anti-tumor immunity in breast cancer patients^46^. These differences underscore the complexity and context dependency of CXCL9/10-CXCR3-mediated T cell regulation. We show that fibroblast-derived Cxcl9/10 promotes lung colonization by directly stimulating the growth of CXCR3^+^ cancer cells. Importantly, we detected CXCR3^+^ subsets of cancer cells also in primary cultures of pleural effusion and ascites samples from patients with metastatic breast cancer where the receptor plays a functional role, indicating a relevance of this direct cellular interaction in human metastasis. Our data showing that CXCL9 and CXCL10 are selectively induced in fibroblasts by highly metastatic cells at early stages of metastasis suggests that they may confer a unique metastatic advantage to cancer cells. Ultimately, interruption of the CXCL9/10-CXCR3-mediated paracrine loop through systemic inhibition of JNK or CXCR3 represents a potentially viable strategy to inhibit metastatic colonization of the lungs in breast cancer patients. Future studies elucidating CXCL9/10-CXCR3-induced signaling may furthermore reveal downstream targets implicated in this crosstalk and may thus shed light on the mechanism by which CXCL9/10 promotes metastatic colonization.

## METHODS

### Cell culture

MDA-MB-231 (MDA, ATCC), MDA-MB-231-LM2 (MDA-LM2, provided by Joan Massagué, RRID:CVCL_5998)^14^, SUM159 (SUM, Asterand Bioscience), SUM159-LM1 (SUM-LM1)^21^ and 4T1 (ATCC) cells were cultured in D10f medium, consisting of DMEM GlutaMAX medium (ThermoFisher Scientific) supplemented with 10 % v/v fetal bovine serum (FBS), 50 U/ml penicillin, 50 μg/ml streptomycin (Sigma-Aldrich) and 1 μg/ml amphotericin B. MRC-5 cells (ATCC) and primary fibroblasts obtained from lungs of healthy NSG or BALB/c mice 6-8 weeks of age were cultured in Minimum Essential Medium Eagle medium with alpha modification (MEMα) (ThermoFisher) supplemented with 1x MEM Non-essential Amino Acid Solution (Sigma-Aldrich), 10 % v/v fetal bovine serum (FBS), 50 U/ml penicillin, 50 μg/ml streptomycin and 1 μg/ml amphotericin B (US Biological).

Primary pleural effusion and ascites samples from metastatic breast cancer patients were cultured in a 1:1 mix of supplemented M199 medium ^47^ and modified M87 medium ^48^ as previously described^21^.

### Pleural effusion and ascites samples

Pleural effusion and ascites samples were obtained from breast cancer patients treated at the National Center for Tumor Diseases Heidelberg (NCT) and the Department of Gynecology at the University Clinic Mannheim. Study approval was obtained from the ethical committees of the University of Heidelberg (case number S-295/2009) and the University of Mannheim (case number 2011-380N-MA) and conformed to the principles of the WMA Declaration of Helsinki and the Department of Health and Human Services Belmont Report. Patients gave written informed consent. Samples were processed as previously described^21, 47^.

### Mouse studies

Animal care and procedures were approved by the governmental review board of the state of Baden-Wuerttemberg, Regierungspraesidium Karlsruhe, under the authorization numbers G-51/13, G-81/16, G-218/16, DKFZ299 and DKFZ356 and followed the German legal regulations. Non-obese diabetic-severe combined immunodeficiency gamma^null^ (NSG, obtained from in-house breeding) and BALB/c (Janvier Labs or Envigo) female mice of 6-8 weeks of age were used for mouse studies. Mice were housed in individually ventilated cages under temperature and humidity control. Cages contained an enriched environment with bedding material.

For lung colonization assays, 10,000 - 200,000 cancer cells were injected in 100 μl PBS via the tail vein (t.v.). Human breast cancer cells and 4T1 mouse mammary tumor cells were previously transduced with a triple reporter expressing the genes herpes simplex virus thymidine kinase 1, green fluorescent proteins (GFP), and firefly luciferase (Fluc)^49^, enabling bioluminescent imaging (BLI) of lung metastatic progression. For BLI, mice were injected intraperitoneally with 150 mg/kg D-luciferin (Biosynth), anesthetized using isoflurane (Orion) and imaged with IVIS Spectrum Xenogen machine (Caliper Life Sciences). Bioluminescent analysis was performed using Living Image software, version 4.4 (Caliper Life Sciences).

For treatment with CXCR3 inhibitor, AMG-487 (Tocris) was reconstituted in DMSO to 8 μg/μl and aliquots were frozen at −20 °C. Prior to injection, aliquots were further diluted in 20 % (2-Hydroxypropyl)-β-cyclodextrin solution, resulting in a final concentration of 2.5 % DMSO. DMSO/Cyclodextrin solution was used as vehicle. Mice received subcutaneous (s.c.) injections with 8 mg/kg AMG-487 every 12 h for the duration of the experiment, starting 12 h before tail vein (t.v.) injection of cancer cells.

To study tumor initiation capacities of CXCR3^+^ 4T1 mouse mammary tumor cells compared to CXCR3^−^, sorted cells were seeded in 10 cm dishes in 10 ml D10f and incubated at 37 °C overnight. The following day, 10,000, 5,000 or 500 CXCR3^+^ or CXCR3^−^ 4T1 cells were mixed with 100,000 BALB/c mouse lung fibroblasts in PBS. Cell suspensions of fibroblasts and CXCR3^+^ or CXCR3^−^ 4T1 cells were implanted subcutaneously (s.c.) into either flanks of BALB/c mice in a 1:1 mix of PBS and Matrigel. Mice were sacrificed after 20 days and tumor formation was recorded. Tumor sizes were measured with a digital caliper and tumor volume was calculated as (width x length x height)/2.

### Fluorescence-activated cell sorting (FACS) and Flow cytometry

For profiling of fibroblasts at different metastatic stages, mice with similar bioluminescence signals from MDA and MDA-LM2 metastasis groups were selected at week 1 and 3 post cancer cell injection. Lungs were digested in 0.5 % (w/v) Collagenase type III (Pan Biotech), 1 % (w/v) Dispase II (Gibco) and 30 μg/ml DNase I in PBS for 30-45 min at 37 °C. Lungs from age-matched healthy mice were used as control group. Single cell suspensions were obtained by pipetting and filtering through 70 μm nylon filters with FACS buffer (2 % FCS in PBS). Cells were pelleted and red blood cells were lysed with ACK buffer (Lonza). Lysis was stopped by FACS buffer and cell pellets were resuspended in PBS and counted using a ViCell Automated Cell Counter. Per 1×10^6^ cells, 100 μl FcR blocking reagent (Miltenyi BioTec, diluted to 1x in FACS buffer) were added and cells were incubated for 10 min on ice. Respective antibody cocktails were prepared in FACS buffer and added 1:1 to cells in FcR blocking reagent. The following antibodies were used at the indicated staining dilutions: CD45-PE (1:3,000, eBioscience), CD11b-PE (1:3,000, BD Biosciences), CD31-PE (1:1,000, eBioscience), CD326(EPCAM)-PE (1:250, eBioscience); CD140a-APC (1:50, eBioscience), CD140b-APC (1:50, eBioscience). Cells were stained for 30 min on ice in the dark. After staining, cells were washed three times in FACS buffer, filtered and resuspended in MEMα medium or FACS buffer containing 3 μg/ml DAPI (BioLegend). Cells were sorted on BD FACSAria1 or FACSAria2 machines and collected in 150 μl Arcturus PicoPure Extraction Buffer. Numbers of fibroblasts per lung were determined by multiplying the total cell count per lung with the percentage of PE-APC+ fibroblasts as analyzed using FlowJo™ V10.

For flow cytometric staining of CXCR3, human breast cancer were detached using Trypsin (when grown as monolayer, Sigma-Aldrich) or pelleted and dissociated using StemPro Accutase (when grown as spheres, Life Technologies), counted and stained with 5 μg PE-conjugated anti-human CD183 (CXCR3) antibody or PE-conjugated mouse IgG1, κ isotype control (BioLegend) per 1 Mio cells in 100 μl FACS buffer for 30 min on ice in the dark. For staining of 4T1 mouse mammary tumor cells, 2.5 μg PE-conjugated anti-mouse CD183 (CXCR3) antibody or Armenian Hamster IgG isotype control per 1 Mio cells in 100 μl FACS buffer were used. After staining, cells were washed three times in FACS buffer and resuspended in FACS buffer containing 3 μg/ml DAPI (BioLegend). Flow cytometric analysis was done on a BD LSRFortessa analyzer and sorting was done on a BD FACSAria1 machine.

### RNA extraction

RNA was extracted from cultured cells with the QIAGEN RNeasy Mini Kit according to the manufacturer’s protocol. RNA from FACS-isolated cells was purified using the Arcturus PicoPure Extraction Kit (ThermoFisher Scientific) according to the manufacturer’s protocol. RNA concentration and purity were measured on a Nanodrop1000 spectrophotometer (Peqlab) or Bioanalyzer 2100 (Agilent).

### RT-qPCR

cDNA was generated from total RNA using the High-Capacity cDNA Reverse Transcription Kit (Applied Biosystems) according to the manufacturer’s protocol. cDNA corresponding to 20-40 ng RNA were used as input for qPCR. Gene expression was analyzed using SYBR Green gene expression assay (Applied Biosystems) on the ViiA 7 Real-Time PCR System (Applied Biosystems) using the following primer pairs: *RPL13A* human housekeeping gene (F: 5’-AGATGGCGGAGGTGCAG-3’, R: 5’-GGCCCAGCAGTACCTGTTTA-3’), human *CXCL9* (F: 5’-GAGTGCAAGGAACCCCAGTAG-3’, R: 5’-GGTGGATAGTCCCTTGGTTGG-3’), human *CXCL10* (F: 5’-TGGCATTCAAGGAGTACCTCTC-3’, R: 5’-GGACAAAATTGGCTTGCAGGA-3’), human *IL1A* (F: 5’-GCTGAAGGAGATGCCTGAGATA-3’, R: 5’-ACAAGTTTGGATGGGCAACTG-3’), human *IL1B* (F: 5’-AACAGGCTGCTCTGGGATTC-3’, R: 5’-AGTCATCCTCATTGCCACTGT-3’), *B2m* murine house-keeping gene (F: 5’-CCTGGTCTTTCTGGTGCTTG-3’, R: 5’-CCGTTCTTCAGCATTTGGAT-3’), murine *Cxcl9* (F: 5’-TCGGACTTCACTCCAACACAG-3’, R: 5’-AGGGTTCCTCGAACTCCACAC-3’), murine *Cxcl10* (F: 5’-GAGAGACATCCCGAGCCAAC-3’, R: 5’-GGGATCCCTTGAGTCCCAC-3’).

### Gene Expression analyses

RNA from sorted fibroblasts was submitted for transcriptomic analysis using Affymetrix GeneChip^®^ Mouse Genome 430 2.0 Arrays after cDNA preamplification using the Ovation Pico WTA System V2 Kit (NuGEN) according to the manufacturer’s protocol. RNA from sorted MDA and MDA-LM2 cancer cells from micrometastatic lungs (1 week post injection) was analyzed using Affymetrix Human Transcriptome Array 2.0 (HTA 2.0) microarrays after cDNA amplification using GeneChip WT Pico Reagent Kit (Affymetrix) according to the manufacturer’s protocol. Raw CEL-files were RMA-normalized, and two-group comparisons were performed using Chipster (version 3.8.0) using empirical Bayes test and Benjamini-Hochberg correction of *P* values. RNA from sorted CXCR3^+^ or CXCR3^−^ SUM-LM1 cells was analyzed using Affymetrix Human U133 Plus 2.0 microarrays after cDNA preamplification using the 3’IVT Pico Reagent Kit (Affymetrix). To generate the CXCR3 signature, PCA clustering was performed on RMA-normalized data and the top 300 genes driving PC1 were retrieved using R (RStudio 1.0.143). The top 65 genes enriched in CXCR3^+^ SUM-LM1 cells from PCA clustering establish the CXCR3 signature.

Gene set enrichment analysis (GSEA) was performed as previously described^50, 51^ and nominal p values were calculated based on random gene set permutations with Benjamini-Hochberg correction. FDR < 0.1 were regarded as statistically significant. Gene Ontology analysis was carried out using Database for Annotation, Visualization and Integrated Discovery (DAVID)^52, 53^. Generated data sets are accessible under the GEO accession number: GSE121947. Additionally, the following gene expression data sets were used: GSE16446 (TOP trial) and GSE14020.

For survival analysis of breast cancer patients, the TOP trial data set was used (GSE16446) as well as data sets from Kaplan-Meier plotter (KM plotter) to analyze basal-like breast cancer samples^54^. Samples were divided into CXCR3S high and low based on median cutoff, using mean expression of the 65 CXCR3S genes. Basal-like breast cancer samples were compiled from the following data sets in KM plotter: For overall survival (Fig. 7G) - GSE1456, GSE16446, GSE16716, GSE20685, GSE20711, GSE3494, GSE37946, GSE42568, GSE45255, GSE7390; for Distant-Metastasis-Free Survival (Supplementary Fig. S8B) - GSE11121, GSE16446, GSE19615, GSE20685, GSE26971, GSE2990, GSE3494, GSE45255, GSE7390, GSE9195; for Relapse-Free Survival (Supplementary Fig. S8C) - E-MTAB-365, GSE11121, GSE12276, GSE1456, GSE16391, GSE16446, GSE16716, GSE17705, GSE19615, GSE2034, GSE20685, GSE20711, GSE21653, GSE2603, GSE2990, GSE31519, GSE3494, GSE37946, GSE42568, GSE45255, GSE4611, GSE5327, GSE7390, GSE9195. ECM gene collections were obtained from the matrisomeproject.edu^55^.

### Oncosphere formation

For 3D cultures, cancer cells were seeded into 75 cm^2^ ultra-low attachment cell culture flasks (Corning) in Onco2 medium, consisting of HuMEC-medium (Invitrogen) supplemented 50 U/ml penicillin (Sigma-Aldrich), 50 μg/ml streptomycin (Sigma-Aldrich), 10 ng/ml basic fibroblast growth factor (bFGF, Invitrogen), 20 ng/ml EGF (Sigma-Aldrich), 5 μg/ml human insulin (Sigma-Aldrich) and 2 % vol/vol B27 (Life Technologies) at a density of 25,000 cells/ml (10 ml per flasks) and incubated at 37 °C for 1 week.

For sphere formation assays, cancer cells were seeded into 96-well ultra-low attachment plates (Corning) in Onco2 medium at a density of 10,000 cells/ml (200 μl per well). 10 μM AMG-487 (Tocris) or vehicle (0.001 % DMSO in Onco2 medium) were added to the medium on day 0 (day of seeding), day 1, 4 and 6. 100 ng/ml recombinant human CXCL9 or CXCL10 (Peprotech) or vehicle (0.1 % BSA in PBS) were added on day 1, 4 and 6. For sphere formation of cancer cells overexpressing CXCL9 and/or CXCL10, cells were seeded as described above without further stimulation. Sphere formation was quantified after 7 days by counting total number of spheres per well using a Zeiss Primovert microscope. Ten wells per condition were quantified and normalized sphere counts were quantified from two to three independent experiments.

### Immunohistochemistry

Mouse lungs were fixed in formalin for 6-8 h at 4 °C, washed and incubated at 4 °C overnight in 30 % Sucrose/PBS. The next day, lungs were washed and embedded in OCT (Sakura). 8 μm sections were cut using a Microm HM 525 cryotome (ThermoFisher Scientific).

For vimentin staining of mouse lungs or αSMA staining of formalin-fixed-paraffin-embedded (FFPE) patient lung metastases, sections were rehydrated with decreasing concentrations of ethanol, quenched with 3 % hydrogen peroxide and antigen retrieval was carried out at 100 °C for 20 min with citrate buffer (pH 6.0, Vector Laboratories) for vimentin staining and pH9 buffer (Vector Laboratories) for αSMA staining. Sections were blocked with 0.1 % BSA containing 0.1 % Triton-X for 2 h at room temperature, followed by incubation with respective primary antibodies (vimentin: Leica Biosystems, clone SRL33, 1:400; αSMA, clone 1A4, 1:100, abcam). Corresponding anti-mouse or anti-rabbit IgG biotinylated secondary antibodies and ABC avidin-biotin-DAB detection kit (Vector laboratories) were used for signal detection according to manufacturer’s instructions. Sections were counterstained with Mayer’s hematoxylin solution (Sigma-Aldrich) for 1 min, dehydrated using increasing concentrations of ethanol and mounted using Cytoseal XYL (ThermoFisher Scientific).

For quantification of sizes of metastatic nodules 1 week and 3 weeks post intravenous injection of MDA and MDA-LM2 cells, vimentin stainings of metastatic lungs (6 mice per group) were analyzed. 7-10 pictures per lung were obtained using 10x objective of an AxioPlan microscope (Carl Zeiss) and longest dimensions of metastatic foci were measured using the Fiji distribution of the ImageJ software (Schindelin et al, 2012).

For hematoxylin and eosin (HE) staining, sections were rehydrated with decreasing ethanol concentrations and stained for 6 min with Mayer’s hematoxylin solution. After washing and short incubation in ethanol supplemented with 0.3 % hydrogen chloride, sections were washed in tap water and counterstained with eosin Y alcoholic solution (Sigma-Aldrich), dehydrated and cleared in xylenes (Sigma-Aldrich) before mounting in Cytoseal XYL (ThermoFisher Scientific). Stainings were analyzed on a Cell Observer motorized widefield microscope (Zeiss).

### Immunofluorescence

Sorted fibroblasts from lungs 1 week post intravenous injection of 200,000 MDA- or MDA-LM2 cells were seeded onto collagen-coated coverslips in 12-well plates in 1 ml MEMα medium. Fibroblasts were cultured for 5 days prior to immunofluorescent staining. For staining, cells were washed with PBS, fixed in 10 % formaldehyde for 30 min on ice, washed and permeabilized with 0.1 % TritonX-100 in PBS-T (PBS containing 0.05 % Tween20) for 10 min at room temperature. Blocking was done in 1 % BSA in PBS for 1 h at room temperature. Primary antibodies against αSMA (abcam) and Ki-67 (eBioscience) were diluted 1:100 in PBST and 200 μl were added per coverslip and incubated for 1 h at room temperature. Afterwards, cells were washed three times with PBS-T for 5 min and incubated with secondary Cy3-coupled goat anti-rat IgG antibody (for Ki-67) or AlexaFluor488-coupled goat anti-mouse IgG (for αSMA) antibodies diluted 1:500 in PBS-T for 1 h at room temperature in the dark. Cells were washed four times and embedded with Prolong Gold Antifade mountant containing DAPI (ThermoFisher Scientific). Pictures were taken on a Cell Observer (Zeiss) using a 10x objective and ZEN software. Percentage of Ki67+ fibroblasts was calculated as follows: (Cy3-positive cells per FOV/DAPI-positive cells per FOV)*100 using the Cell Counter Plugin in ImageJ. Dots represent 3 biological replicates, for which 2-3 fields of view were quantified each. For αSMA quantification, mean αSMA intensities (gray values) of 20 sorted fibroblasts were measured using ImageJ. Dots represent average gray value per biological replicate.

### Production of conditioned medium

For production of conditioned medium (CM), cancer cells were seeded in 10 cm culture dishes in 10 ml D10f medium to 70-80 % confluency. Cells were washed once in PBS and 6 ml serum-free MEMα medium were added. After incubation at 37 °C for 48 h, medium was aspirated and filtered through a 0.45 μm filter. Conditioned medium was used directly or aliquoted and frozen at −80 °C.

### Stimulation of fibroblasts

For treatment of fibroblasts with recombinant IL-1α/β or cancer cell CM, 1.5 - 2 x10^5^ MRC-5 cells or primary mouse lung fibroblasts obtained by culturing in fibroblast-specific MEMα medium from NSG mice, IL1R-KO Il1r1^tm1Roml^ C57BL/6 mice or C57BL/6 wildtype mice were seeded in 1 ml MEMα medium per 6-well. When treatments included IL1R blockade or NF-κB inhibition, MRC-5 were seeded in the presence of 20 μg/ml anti-IL-1R1 antibody or Normal Goat IgG Control (R&D), or 5 μM JSH-23 (Sigma-Aldrich) or 0.1 % DMSO as a vehicle. The next day, medium was aspirated, fibroblasts were washed with 1 ml PBS per well, and 1 ml CM or serum-free MEMα were added. 1 ng/ml human recombinant IL-1α/β (Peprotech) or 0.1 % BSA in PBS as carrier control as well as 20 μg/ml anti-IL-1R1/IgG control or 5 μM JSH-23/ 0.1 % DMSO vehicle were added to the respective wells. After 48 h incubation at 37 °C, fibroblasts were washed with PBS and lysed in RLT buffer (Qiagen) for RNA extraction.

### Gel contraction assay

NSG mouse lung fibroblasts were adjusted to 1.5 x 10^5^ cells/ml in PBS. 800 μl of the fibroblast suspension were pipetted into a 1.5 ml Eppendorf tube and 400 μl of a 3 mg/ml bovine collagen solution (Advanced BioMatrix) were added to the cell suspension and mixed by pipetting. 8 μl of 1 M NaOH were added to the cell-collagen mixture and the solution was mixed by pipetting. 1 ml of the mixture was immediately transferred to a 12-well plate and gels were allowed to solidify at room temperature for 20 min. Cancer cell CM obtained from 250,000 MDA or MDA-LM2 cancer cells seeded in 2 ml MEMα medium per 6-well for 48 h was filtered through 0.45 μm filter and 1 ml CM or MEMα control medium was added per well. Gels were dissociated from the wells by gently running the tip of a 200 μl pipet tip along gel edges and swirling the plate. 12-well plates were incubated at 37 °C and gel diameters were recorded after 24 h.

### JNK inhibition

MDA-MB-231-LM2 cells were seeded at a density of 250,000 cells/6-well or 1×10^6^ cells/10 cm dish in D10f medium containing 5 μM CC-401 (Santa Cruz Biotechnology) or 0.1 % DMSO as a vehicle. The following day, medium was exchanged and cells were stimulated with fresh D10f containing 5 μM CC-401 or 0.1% DMSO as a vehicle. After 48 h, medium was aspirated, cells were washed in PBS and lysed in RLT buffer (Qiagen) for RNA extraction or counted for injection.

### Knockdown of *IL1A/B* in cancer cells

*IL1A/B* double knockdown was generated in MDA-LM2 cells with miR-E lentiviral vectors^56^ expressing shRNA against *IL1A/B* gene products. miR-E *IL1A* hairpins were produced from the StagBFPEP lentiviral vector, a modified version of the original SGEP vector kindly provided by Johannes Zuber (IMP-Research Institute of Molecular Pathology GmbH, Vienna), in which the constitutively expressed GFP protein was replaced by the tagBFP protein. miR-E *IL1B* hairpins were produced from the StdTomatoEZ lentiviral vector, a modified version of the original SGEP vector, in which the constitutively expressed GFP protein was replaced by the tdTomato protein and the Puromycin resistance cassette was replaced by the Zeocin resistance cassette. miR-E shIL1A/B oligonucleotides were designed using the shERWOOD algorithm^57^. The following hairpins were used: shIL1A: 5’-TGCTGTTGACAGTGAGCGCCCTGAGCAATGTGAAATACAATAGTGAAGCCACAGATGTATTGTATTTCACATTGCTCAGGATGCCTACTGCCTCGGA-3’, shIL1B: 5’-TGCTGTTGACAGTGAGCGCCAATAACAAGCTGGAATTTGATAGTGAAGCCACAGATGTATCAAATTCCAGCTTGTTATTGATGCCTACTGCCTCGGA-3’. Oligonucleotides were amplified by PCR using the Q5 High-Fidelity DNA Polymerase (New England Biolabs) and the following primers: miRE-Xho-fw: 5’-TGAACTCGAGAAGGTATATTGCTGTTGACAGTGAGCG-3’, miRE-EcoOligo-rev: 5’-TCTCGAATTCTAGCCCCTTGAAGTCCGAGGCAGTAGGC-3’. PCR products containing shIL1A, shIL1B and non-silencing miR-Es were subcloned into the StagBFPEP and StdTomatoEZ recipient vectors via EcoRI-HF and XhoI restriction sites. Lentiviral particles were produced by transfecting HEK293T cells with StagBFPEP-miR-E shIL1A, StdTomatoEZ shIL1B, StagBFPEP-miR-E shControl or StdTomatoEZ shControl together with pMD2G and psPAX2 packaging plasmids using Lipofectamine 2000 (Invitrogen). Supernatants containing lentiviral particles were used to infect MDA-LM2 cancer cells overnight in the presence of 8 μg/ml polybrene (Sigma-Aldrich). Infected cells were selected with 2 μg/ml puromycin (Invitrogen) or 0.4 mg/ml zeocin (ThermoFisher Scientific) in D10f medium for 3-7 days until uninfected control cells were dead. MDA-LM2 cells were first infected with shIL1B/shControl StdTomatoEZ lentiviral particles and selected, and subsequently infected with shIL1A/shControl StagBFPEP lentiviral particles for double knockdown.

### Overexpression of *CXCL9/10* in cancer cells

Human *CXCL9* and *CXCL10* cDNA flanked by XhoI and BamHI restriction sites were ordered as GeneArt™ Strings™ DNA Fragments (Invitrogen). DNA strings were digested with XhoI and BamHI in CutSmart buffer (New England Biolabs) for 1 h at 37 °C and purified using QIAquick PCR Purification Kit (Qiagen). The pLVX-Puro lentiviral expression vector (Clontech) with constitutive CMV promoter was used to insert *CXCL9* or *CXCL10* cDNA. For combined overexpression of *CXCL9* and *CXCL10*, a pLVX-Hygro lentiviral vector was additionally generated. For this purpose, the pLVX-FADD-DD plasmid was obtained from Addgene, which was a gift from Joan Massagué (Addgene plasmid # 58263)^18^ and contains the pLVX-IRES-Hygro backbone (Clontech). The FADD-DD insert was replaced by subcloning the multiple cloning site of the pLVX-Puro backbone plasmid via SnaBI and BamHI restriction. pLVX-Puro and pLVX-Hygro lentiviral vectors were then digested with XhoI and BamHI in CutSmart buffer for 1 h at 37 °C, purified using QIAquick Gel Extraction Kit (Qiagen) and dephosphorylated using Antarctic phosphatase (New England Biolabs). After ligation of insert and vector DNA with T4 DNA ligase (New England Biolabs) for 30 min at room temperature, followed by 65 °C heat inactivation for 5 min, ElectroMAX Stbl4 Competent bacterial cells (Life Technologies) were transformed by electroporation and DNA sequencing was used to confirm correct insertion of *CXCL9/10* sequences. To produce lentiviral particles, HEK293T cells were cotransfected with pLVX-Puro or pLVX-Hygro vectors containing *CXCL9* or *CXCL10* inserts and the packaging plasmids psPAX2 and pMD2G using Lipofectamine 2000 (Invitrogen). Supernatant containing lentiviral particles was used to infect cancer cells in the presence of 8 μg/ml polybrene (Sigma-Aldrich) overnight. Infected cells were selected with 2 μg/ml puromycin (Invitrogen) or Hygromycin B (Life Technologies) for 3-7 days, until uninfected control cells were dead and overexpression of *CXCL9/10* was confirmed by RT-qPCR.

### Enzyme-linked immunosorbent assay (ELISA)

For detection of human IL-1α and IL-1β and mouse Cxcl9 and Cxcl10 in metastatic lungs, whole lungs from mice bearing MDA-LM2 macrometastases (3 weeks post tail vein injection) were harvested and lysed in 1 ml Cell Lysis Buffer 2 (R&D) using the gentleMACS Dissociator (Miltenyi Biotec). As a control, lungs from healthy age-matched mice were lysed. For detection of human IL-1α and IL-1β, lung homogenates were diluted 1:1 in Calibrator Diluent RD6C. For detection of mouse Cxcl9 and Cxcl10, lung homogenates were diluted 1:10 in Cell Lysis Buffer 2 (R&D). Quantikine Human IL-1α ELISA Kit (R&D), Human IL-1β/IL-1F2 Quantikine ELISA Kit (R&D), Mouse Cxcl10 DuoSet ELISA (R&D) and Mouse Cxcl9 Quantikine ELISA (R&D) were used according to the manufacturer’s protocol. Standard curves were calculated in GraphPad Prism version 7.02 by linear regression. To account for initial dilution of lung homogenates, interpolated IL-1α/β concentrations need to be multiplied by 2 and interpolated Cxcl9/10 concentrations need to be multiplied by 10 to determine final protein concentrations per lung.

### Chromatin immunoprecipitation (ChIP)

We performed chromatin immunoprecipitation (ChIP) on 4-5×10^6^ MDA-LM2 or SUM-LM1 cells using the PierceTM Magnetic ChIP Kit (ThermoFisher Scientific) according to manufacturer’s instructions with 10 μg rabbit IgG isotype control or c-Jun antibody (Cell Signaling). Primers for ChIP-qPCR were designed flanking known consensus and tracked AP-1/c-Jun binding sites in proximity to IL1A and IL1B promoter regions, as assessed by the USCS Genome Browser^58^. SYBR green (Applied Biosystems) qPCR was conducted using the following primer sequences: *IL1A* (Primer pair 1: F: 5’-GGCTGTAGCTTTAGAGAAGGCA-3’, R: 5’-GGCGTTTGAGTCAGCAAAGG-3’, Primer pair 2: F: 5’-CCTTTGCTGACTCAAACGCC-3’, R: 5’-AGCCACGCCTACTTAAGACAA-3’), *IL1B* (Primer pair 1: F: 5’-CCTTGTGCCTCGAAGAGGTT-3’, R: 5’-TCTCAGCCTCCTACTTCTGCT-3’, Primer pair 2: F: 5’-ATGGGTACAATGAAGGGCCAA-3’, R: 5’-GCTCCTGAGGCAGAGAACAG-3’). qPCR was analyzed with the Viia 7 Real-Time PCR System (Applied Biosystems).

### Statistical Analysis

Statistical analyses were performed as described in figure legends. *P* values ≤ 0.05 were considered statistically significant and statistical tests were two-tailed unless otherwise indicated. All functional *in vivo* experiments were based on extensive *in vitro* results that suggested one-directional effects, and thus one-tailed t-tests were used when analyzing tumor burden and metastasis in mice. Ratio-paired one-tailed t-tests were used to analyze changes in *CXCL9/10* expression in cultured fibroblasts upon combination of a stimulus (CM or recombinant cytokines) with a blocking treatment (NF-κBi, IL1Rab, *IL1A/B* knockdown) to account for differences in baseline fold change compared to control group. Bars represent mean and error bars represent standard deviation (SD). For Kaplan-Meier analyses of breast cancer patients (GSE16446), statistical differences in survival curves were calculated by log-rank (Mantel–Cox) test. Patients were classified as CXCR3S high or low based on mean normalized expression of the 65 genes comprising the CXCR3 signature, with median cutoff. The GSE14020 gene expression data set was used to study the correlation between *CXCL9* and *CXCL10* expression as well as mean expression of *CXCL9/10* and *IL1A/B*. Maximum probe sets were analyzed: *CXCL9*: 203915_at, CXCL10: 204533_at, *IL1A*: 208200_at, *IL1B*: 205067_at. Gene expression values for each gene or gene pair within each sample were associated by linear regression using Pearson correlation coefficient (r). Statistical analyses of microarray data generated in this study were calculated with Chipster. Statistical analyses of Gene Ontology was calculated DAVID^52, 53^, and statistical analysis of enrichment of gene sets was conducted using GSEA^50, 51^. For GSEA, an FDR<0.1 was considered statistically significant. All other statistical analyses were performed using GraphPad Prism version 7.02 for Windows.

## ACKNOWLEDGEMENTS

We would like to thank S. Acharyya, A.K. Biswas, S. Vanharanta and M. Bettess for critical reading of the manuscript and helpful discussion. We thank J. Massagué for sharing breast cancer cell lines. We are grateful to the microarray unit of the DKFZ Genomics and Proteomics Core Facility for providing the Affymetrix Gene Expression Arrays and related services. Moreover, we would like to thank M. Socher, P. Prückl and the DKFZ Central Animal Laboratory for advice on animal protocols and husbandry and DFKZ Flow Cytometry Core Facility. We also thank M. Brom from the Light Microscopy unit of the DKFZ Imaging and Cytometry Core Facility for technical assistance. The authors are also indebted to K. Decker, A. Riedel and A. Krüger for sharing experimental reagents and for other support. M.P. was supported by a scholarship from the Helmholtz International Graduate School for Cancer Research. T.H. has been funded by the International Tenure Track program of University of Tsukuba. This work was funded by the Dietmar Hopp Foundation.

## AUTHOR CONTRIBUTIONS

M.P. and T.O. designed experiments, analyzed data and wrote the manuscript. M.P. performed *in vitro* and *in vivo* experiments and computational analyses. J.I-R. carried out ChIP, helped with animal experiments and *in vitro* work. J.M. helped with animal experiments and molecular cloning, T.H. helped with processing and sorting of mouse lungs and L.W. helped with *in vitro* experiments. M.A.G.E. contributed to experiments involving IL1R-KO mice. H-P.S. provided metastasis samples from breast cancer patients and advised on pathology. S.S., M.S. and A.S. provided clinical advice and oversaw collection of patient samples. J.I-R., T.H. and A.T. contributed to experimental design and manuscript editing. T.O. supervised the research. All authors read and discussed the manuscript.

---

**Supplementary Figure 1.**
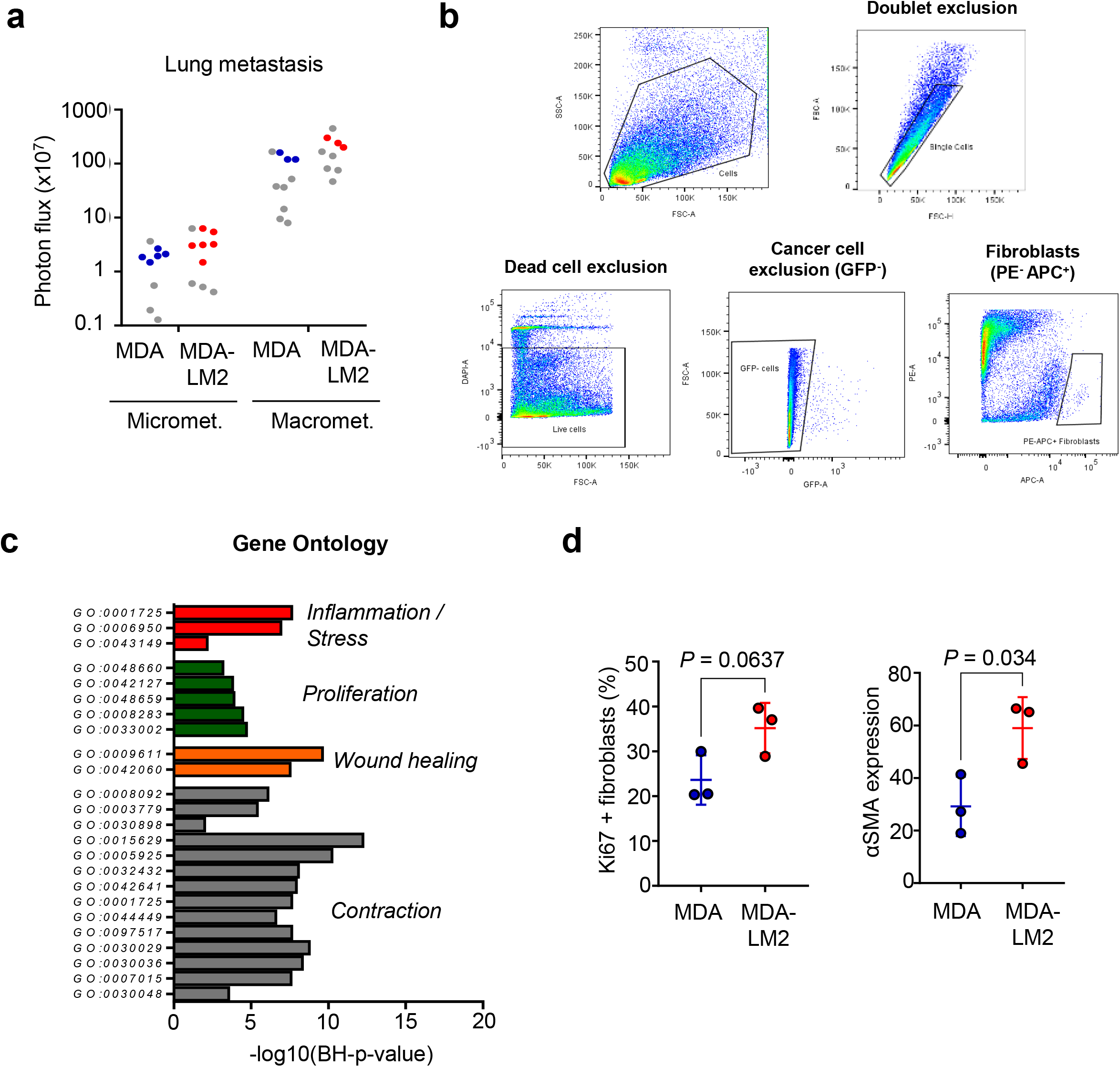
**a**, Lung bioluminescence in mice harboring MDA and MDA-LM2 micro- and macrometastases. Mice used for fibroblast isolation and transcriptomic screen are labelled in blue (MDA) or red (MDA-LM2). Each point represents an individual mouse. **b**, FACS strategy used to isolate fibroblasts from lung metastases. Dead cells were excluded by DAPI stain, cancer cells were excluded by GFP expression, fibroblasts were further enriched by excluding CD45-, CD11b-, EPCAM- and CD31-expressing cells and gating on CD140a- or CD140b-expressing cells. **c**, Gene ontology analysis of top upregulated genes accounting for shift in PCA in fibroblasts from MDA-versus MDA-LM2-micrometastases as shown in Figure 1E. **d**, Quantification of Ki67 and αSMA expression in fibroblasts from Figure 1g. *P* values were calculated by unpaired two-tailed t-tests.

**Supplementary Figure 2.**
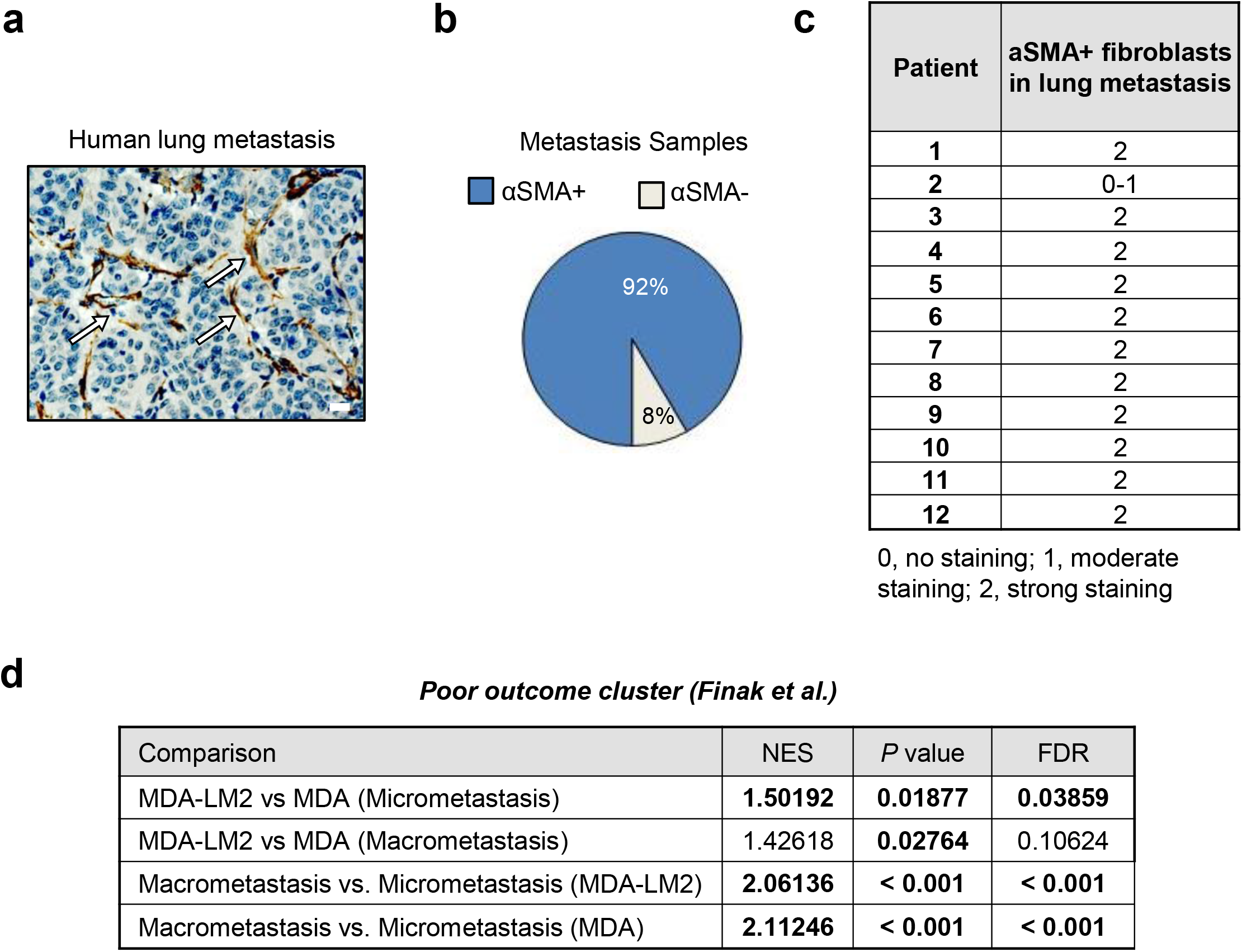
**a-c**, Immunohistochemical analysis of αSMA in human breast cancer metastases in lungs. Representative example of αSMA-expressing fibroblasts (a, white arrows). Scale bar, 20 μm. αSMA expression in lung metastases was analyzed from 12 breast cancer patients (b,c). **d**, GSEA of poor outcome stromal signature (60) in MDA or MDA-LM2 associated MAFs in lungs harboring micro- or macrometastasis. NES, normalized enrichment score. FDR, false discovery rate. *P* values were determined by random permutation tests.

**Supplementary Figure 3.**
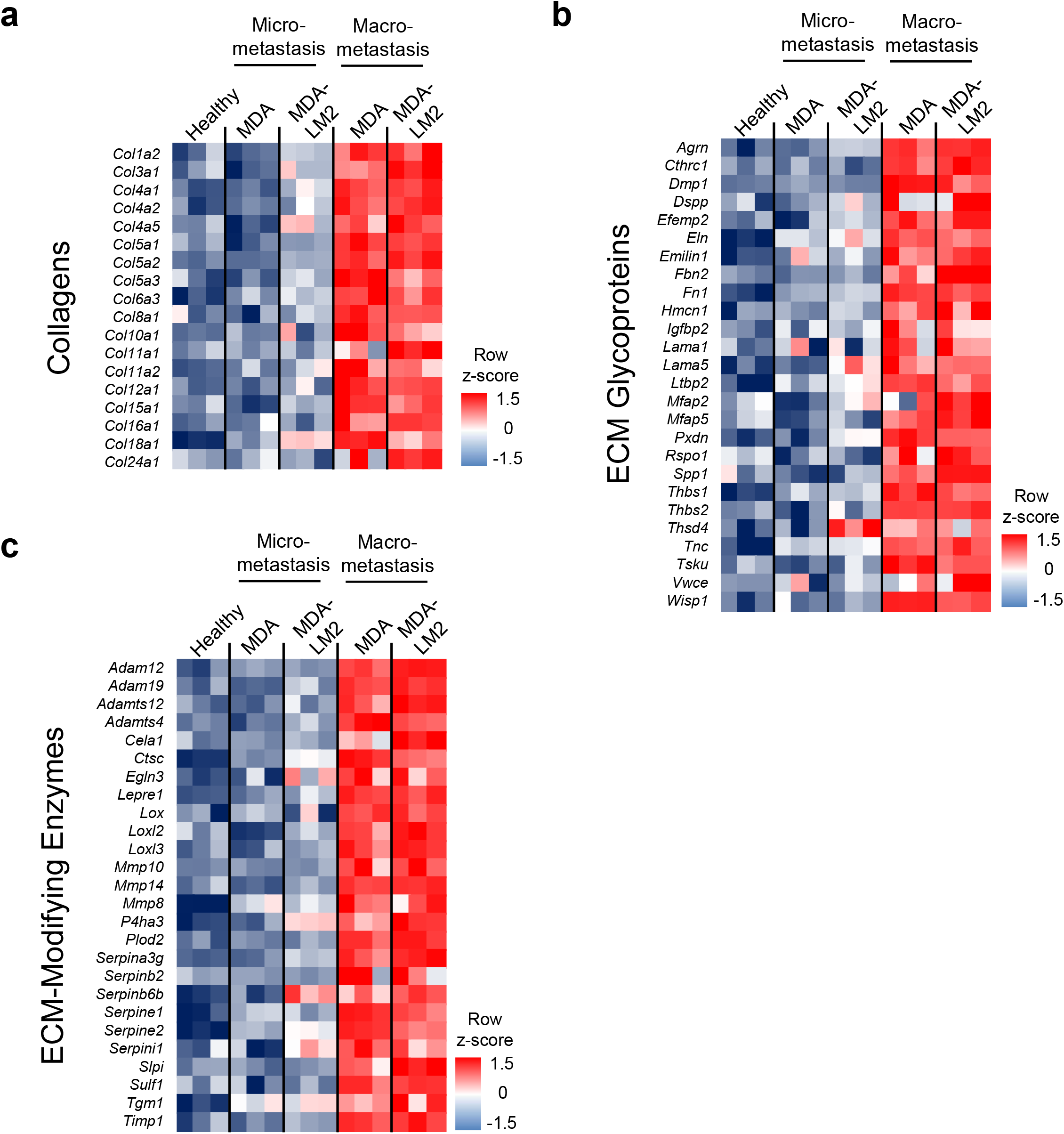
**a-c**, Heatmaps of normalized gene expression of genes encoding collagens (a), ECM glycoproteins (b) and ECM-modifying enzymes (c) in isolated fibroblasts of healthy, micrometastatic and macrometastatic lungs. Gene collections were obtained from the matrisomeproject.mit.edu (55).

**Supplementary Figure 4.**
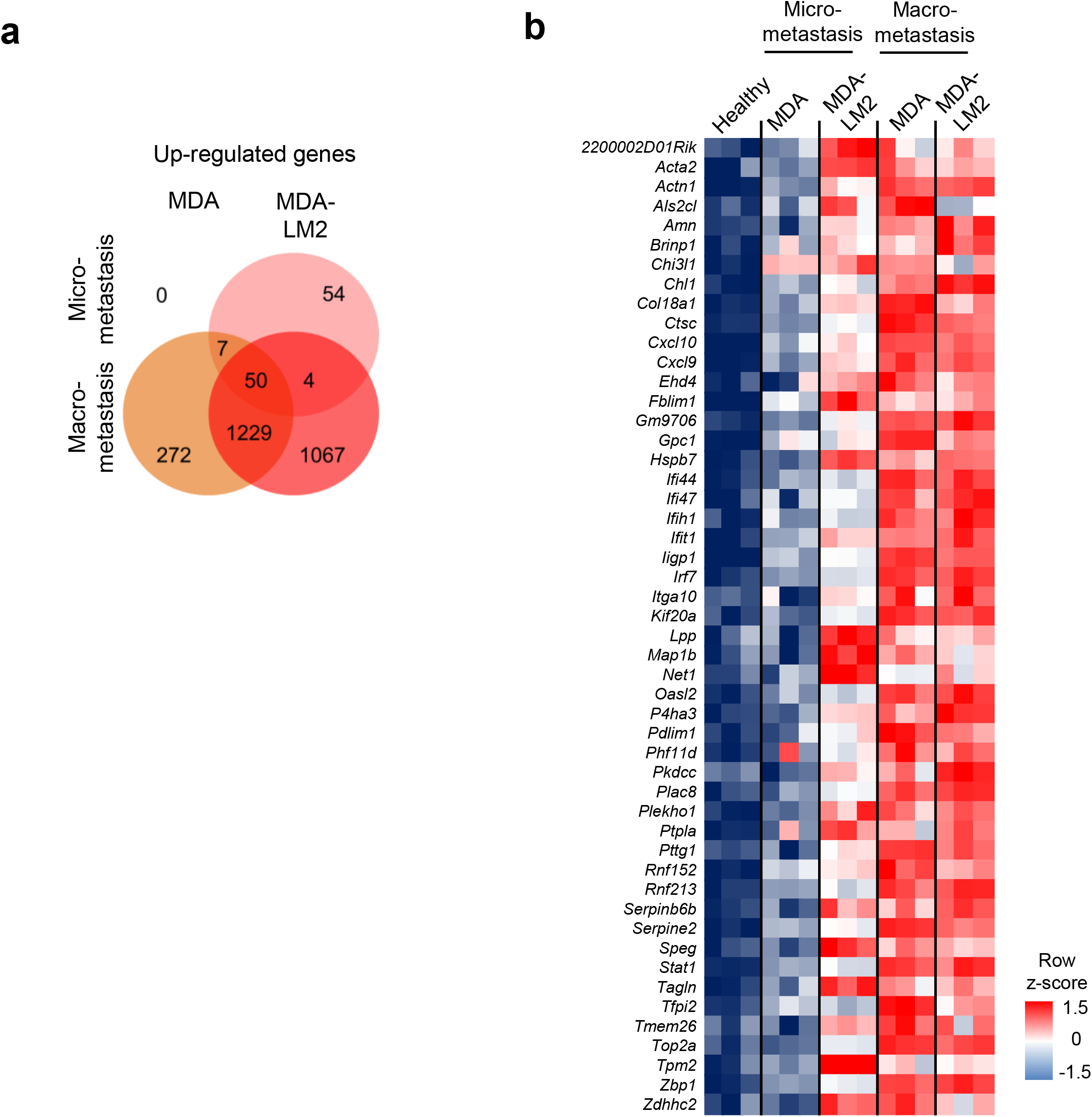
**a**, Venn diagram indicating the number of genes significantly induced (BH-P value < 0.05) in fibroblasts isolated from lungs harboring MDA and MDA-LM2 micro- and macrometastasis. **b**, Heatmap of normalized gene expression from fibroblasts from healthy, MDA and MDA-LM2 micro- and macrometastatic lungs. Shown are genes significantly induced (BH-*P* value < 0.05) in fibroblasts from lungs with MDA-LM2 micrometastasis that overlap with induction at macrometastatic stage.

**Supplementary Figure 5.**
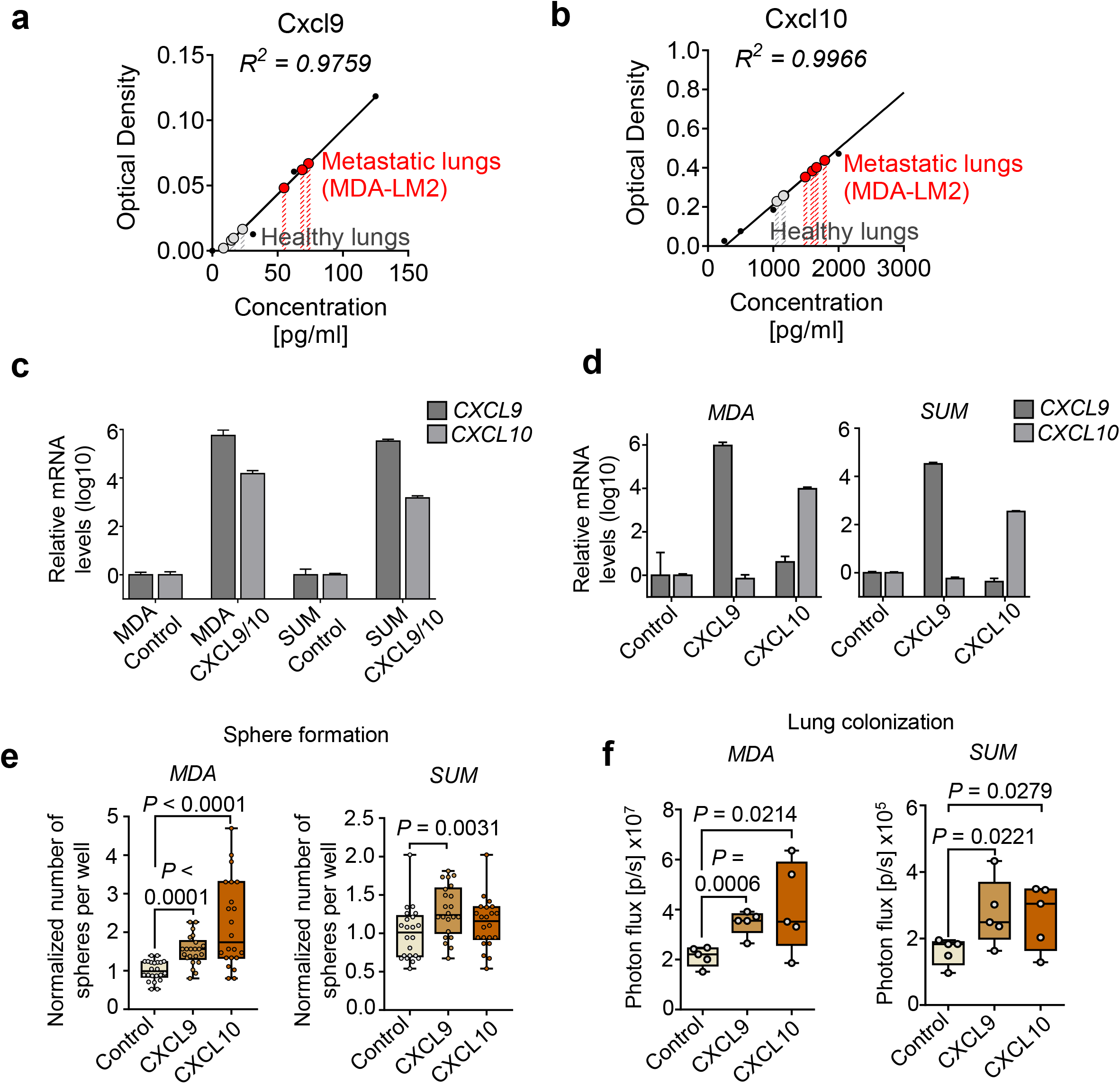
**a-b**, Cxcl9 and Cxcl10 ELISAs from lung homogenates of n=4 healthy mice and n=4 mice harboring MDA-LM2 macrometastases (week 3). Samples were diluted 1:10, thus final concentration per lung is 10-fold as high. **c-d**, *CXCL9/10* mRNA levels in MDA and SUM breast cancer cells upon overexpressing *CXCL9* and *CXCL10* together (c) or individually (d). Expression was determined by RT-qPCR. **e**, Formation of oncospheres by MDA or SUM breast cancer cells overexpressing *CXCL9* or *CXCL10* versus a vector control. Sphere numbers per well were normalized to the average number in the control group. Data represent two independent experiments with quantification of 10 and 12 wells per condition, respectively. **f**, Lung colonization by MDA or SUM breast cancer cells upon ectopic expression of *CXCL9* or *CXCL10* or a vector control, measured using bioluminescence. For (e) and (f), Boxes depict median with upper and lower quartiles. Whiskers show minimum to maximum and dots indicate each individual data point. *P* values were determined by unpaired one-tailed t-tests.

**Supplementary Figure 6.**
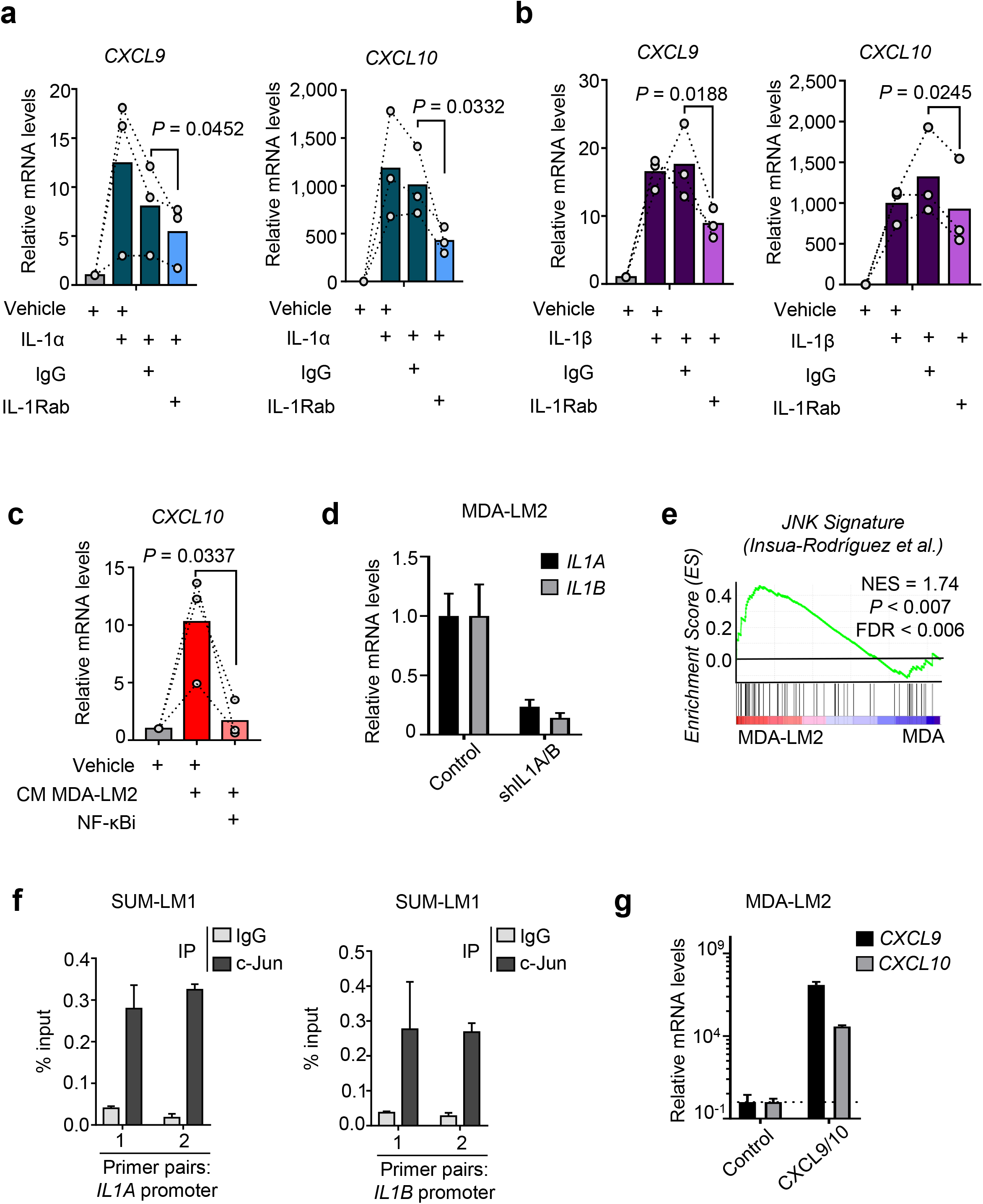
**a-b**, *CXCL9/10* expression in MRC-5 fibroblasts treated with 1 ng/mL recombinant IL-1α (a) or IL-1β (b) in combination with 20 μg/ml IL1R neutralizing antibody or IgG isotype control for 48 h. Expression was analyzed by RT-qPCR. *P* values were calculated by ratio-paired one-tailed t-tests; *n* = 3 independent experiments. **c**, CXCL10 expression in fibroblasts treated with conditioned media (CM) from MDA-LM2 breast cancer cells alone or in combination with 5 μM NF-κB inhibitor (NF-κBi). *P* value was determined by ratio-paired one-tailed t-test; *n* = 3 independent experiments. **d**, *IL1A* and *IL1B* expression determined by RT-qPCR in control and double *IL1A/B* knockdown MDA-LM2 cancer cells. **e**, Enrichment of a JNK response signature (Insua-Rodriguez et al., 2018) in cultured MDA-LM2 versus MDA parental breast cancer cells (14). NES, normalized enrichment score. *P* value was determined by random permutation test. **f**, CHIP-qPCR analysis of c-Jun binding to *IL1A* and *IL1B* promoter chromatin in SUM-LM1 cells. **g**, Ectopic *CXCL9/10* expression in MDA-LM2 breast cancer cells analyzed by RT-qPCR.

**Supplementary Figure 7.**
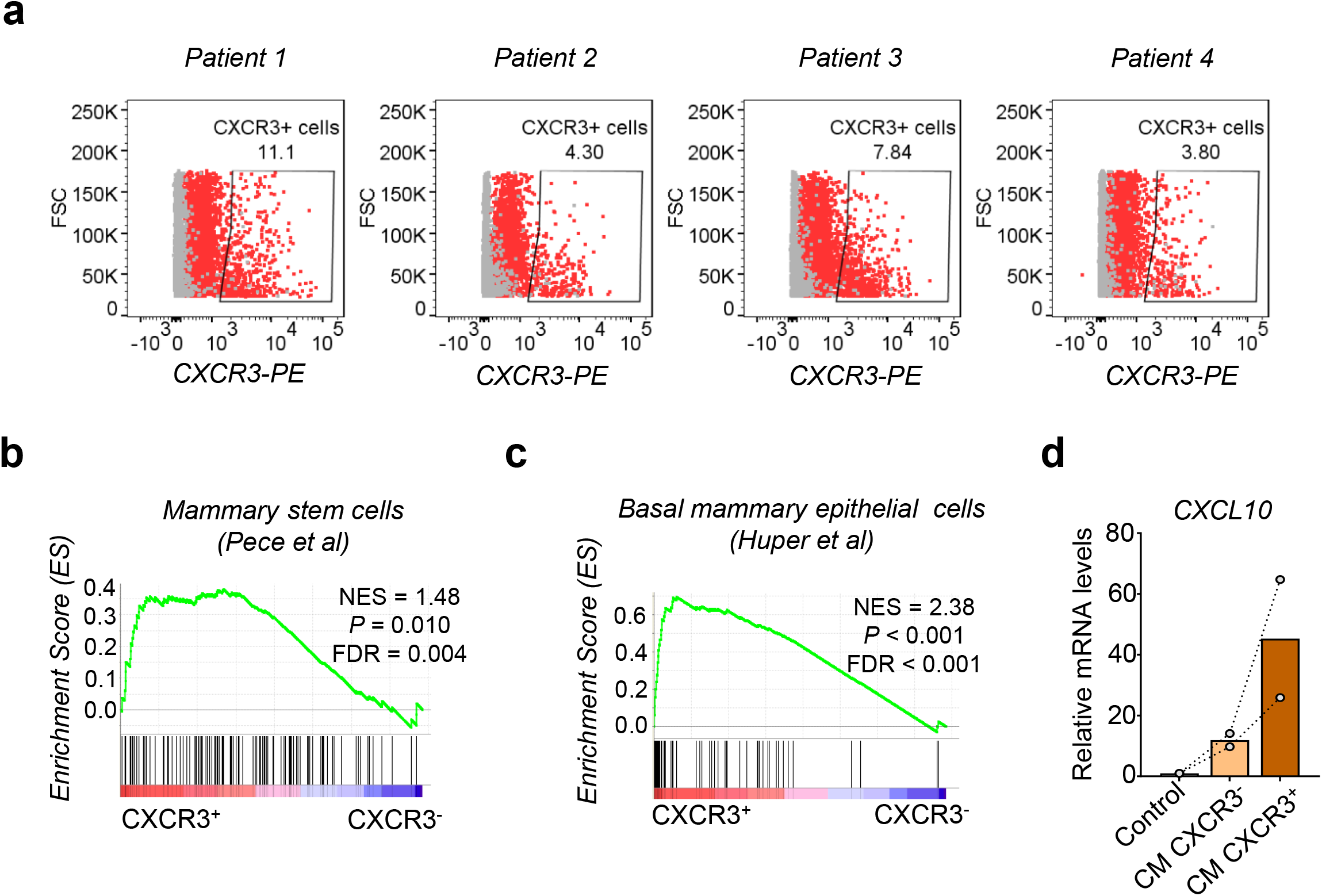
**a**, CXCR3-positive (CXCR3^+^) subpopulations in primary human metastatic cells isolated from pleural effusions (patients 1 and 4) or ascites (patients 2 and 3) of breast cancer patients as determined by flow cytometry. CXCR3-PE staining in red, isotype in grey. Percentages of CXCR3^+^ cells are shown. **b-c**, Enrichment of indicated gene sets in CXCR3^+^ compared to CXCR3^−^ SUM-LM1 breast cancer cells. NES, normalized enrichment score. *P* values were determined by random permutation tests. **d**, *CXCL10* expression in MRC-5 human lung fibroblasts treated with control medium or conditioned medium (CM) from CXCR3^+^ and CXCR3^−^ 4T1 mammary tumor cells for 48 h. Two independent experiments are shown.

**Supplementary Figure 8.**
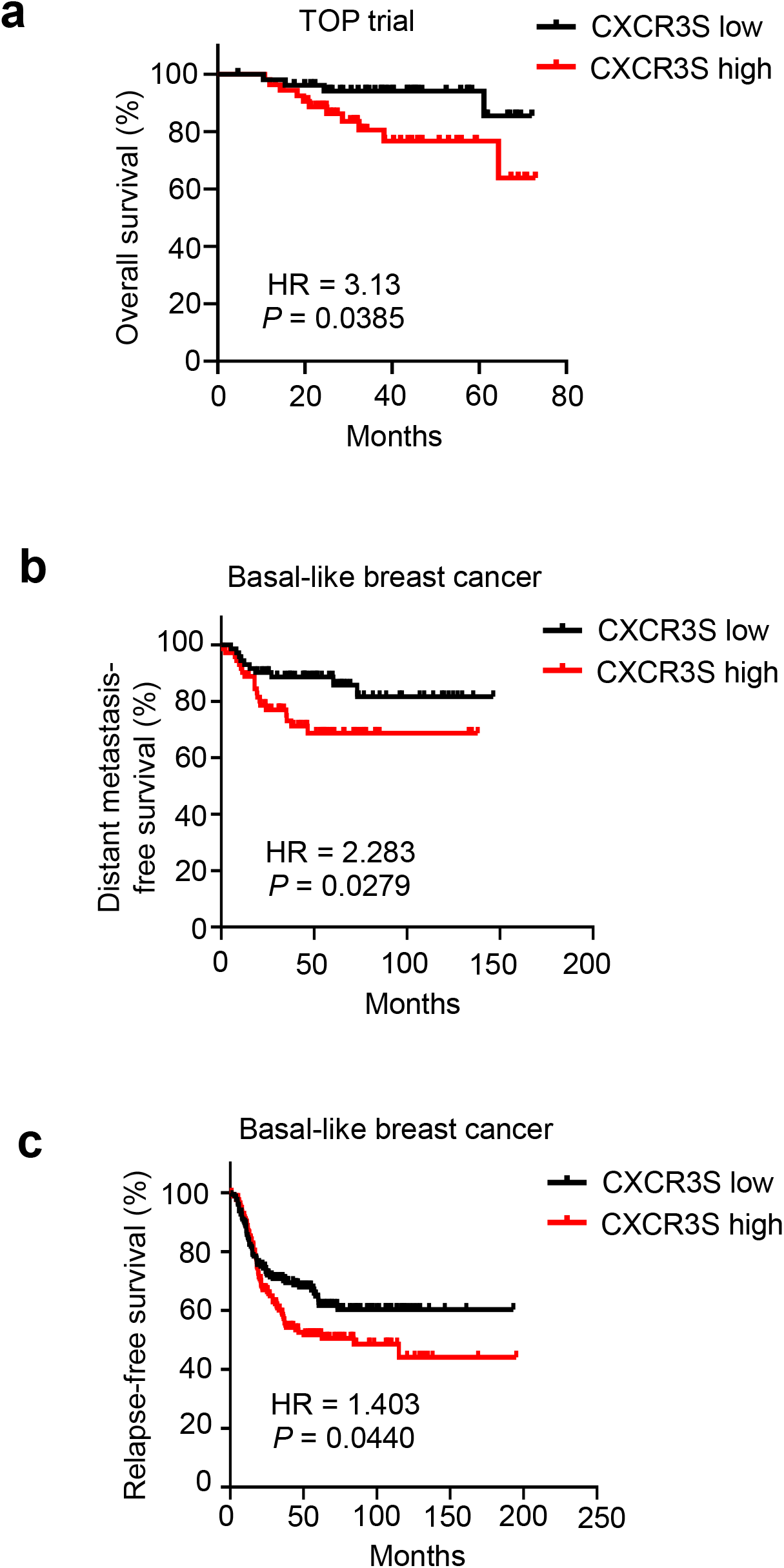
**a-c**, Kaplan-Meier analyses of breast cancer patients, associating CXCR3^+^ cell signature (CXCR3S, mean expression of 65 genes) with overall survival (a, TOP trial data set, *n* = 107 patients), distant metastasis-free survival (b, compiled data set from basal-like breast cancer, KM plotter, *n* = 145) or relapse-free survival (c, compiled data set from basal-like breast cancer, KM plotter, *n* = 360). Median cutoff was used to group patients into CXCR3S low and high. HR, hazard ratio. *P* values were determined by log-rank tests.

**Supplementary Table 1.** GSEA on gene expression profiles of fibroblasts isolated from lungs of mice with growing MDA231 (MDA) and MDA231-LM2 (MDA-LM2) metastases. Shown are normalized enrichment scores (NES) of fibroblasts isolated at 1 week or 3 weeks post injection. False discovery rate (FDR) < 0.1.

| Cellular Responses | NES Micrometastasis |  | NES Macrometastasis |  |
| --- | --- | --- | --- | --- |
|  | MDA | MDA-LM2 | MDA | MDA-LM2 |
| BENPORATH_CYCLING_GENES |  |  | 2.369 | 2.252 |
| BENPORATH_PROLIFERATION |  |  | 2.466 | 2.336 |
| BIOCARTA_CELLCYCLE_PATHWAY |  |  | 1.785 | 1.868 |
| BIOCARTA_G1_PATHWAY |  |  | 1.903 | 2.032 |
| BIOCARTA_G2_PATHWAY |  |  | 1.776 | 1.736 |
| CHANG_CYCLING_GENES |  | 2.064 | 2.959 | 2.903 |
| CHIANG_LIVER_CANCER_SUBCLASS_PROLIFERATION_UP |  | 1.708 | 2.898 | 2.749 |
| KAUFFMANN_DNA_REPLICATION_GENES |  |  | 1.893 | 1.884 |
| KEGG_CELL_CYCLE |  |  | 2.204 | 2.252 |
| KEGG_DNA_REPLICATION |  | 1.880 | 1.868 | 2.102 |
| REACTOME_CELL_CYCLE |  |  | 2.373 | 2.430 |
| REACTOME_CELL_CYCLE_CHECKPOINTS |  | 1.715 | 2.226 | 2.417 |
| REACTOME_CELL_CYCLE_MITOTIC |  |  | 2.447 | 2.506 |
| REACTOME_CYCLIN_E_ASSOCIATED_EVENTS_DURING_G1_S_TRANSITION |  | 1.921 | 2.186 | 2.262 |
| REACTOME_DNA_REPLICATION |  | 1.610 | 2.391 | 2.504 |
| REACTOME_DNA_STRAND_ELONGATION |  | 1.889 | 2.018 | 2.085 |
| REACTOME_G0_AND_EARLY_G1 |  |  | 1.743 | 1.668 |
| REACTOME_G1_PHASE |  |  | 1.866 | 1.831 |
| REACTOME_G1_S_TRANSITION |  | 1.973 | 2.373 | 2.421 |
| REACTOME_LAGGING_STRAND_SYNTHESIS |  | 1.668 | 1.660 | 1.748 |
| REACTOME_M_G1_TRANSITION |  | 2.121 | 2.305 | 2.402 |
| REACTOME_MITOTIC_G1_G1_S_PHASES |  | 1.834 | 2.429 | 2.515 |
| REACTOME_MITOTIC_G2_G2_M_PHASES |  |  | 1.552 | 1.615 |
| REACTOME_MITOTIC_M_M_G1_PHASES |  |  | 2.323 | 2.430 |
| REACTOME_MITOTIC_PROMETAPHASE |  |  | 2.184 | 2.260 |
| REACTOME_REGULATION_OF_MITOTIC_CELL_CYCLE |  | 1.942 | 2.375 | 2.451 |
| REACTOME_S_PHASE |  | 2.122 | 2.399 | 2.500 |
| REACTOME_SYNTHESIS_OF_DNA |  | 2.186 | 2.347 | 2.427 |
| ROSTY_CERVICAL_CANCER_PROLIFERATION_CLUSTER |  | 1.953 | 3.137 | 3.106 |
| WHITFIELD_CELL_CYCLE_G1_S |  |  | 1.694 | 1.535 |
| WHITFIELD_CELL_CYCLE_G2 |  |  | 2.164 | 2.050 |
| WHITFIELD_CELL_CYCLE_G2_M |  |  | 2.407 | 2.344 |
| WHITFIELD_CELL_CYCLE_LITERATURE |  | 1.624 | 2.584 | 2.606 |
| WHITFIELD_CELL_CYCLE_M_G1 |  |  | 1.831 | 1.516 |
| WHITFIELD_CELL_CYCLE_S |  |  | 1.647 | 1.594 |
| ZHANG_PROLIFERATING_VS_QUIESCENT |  |  | 1.904 | 1.900 |
| PLASARI_TGFB1_SIGNALING_VIA_NFIC_10HR_UP |  | 1.980 | 1.429 | 1.573 |
| PLASARI_TGFB1_SIGNALING_VIA_NFIC_1HR_UP |  |  | 1.543 | 1.630 |
| PLASARI_TGFB1_TARGETS_10HR_UP |  | 1.672 | 2.786 | 2.662 |
| PLASARI_TGFB1_TARGETS_1HR_UP |  |  | 2.347 | 2.112 |
| REACTOME_SIGNALING_BY_TGF_BETA_RECEPTOR_COMPLEX |  |  | 1.737 | 1.657 |
| VERRECCHIA_RESPONSE_TO_TGFB1_C1 |  |  | 1.501 |  |
| VERRECCHIA_RESPONSE_TO_TGFB1_C5 |  | 1.758 | 1.501 | 1.742 |
| VERRECCHIA_DELAYED_RESPONSE_TO_TGFB1 |  |  | 1.767 | 1.814 |
| VERRECCHIA_EARLY_RESPONSE_TO_TGFB1 |  |  | 1.569 | 1.467 |
| SEKI_INFLAMMATORY_RESPONSE_LPS_UP |  |  | 2.934 | 2.690 |
| BIOCARTA_INFLAM_PATHWAY |  |  | 1.701 | 1.593 |
| OKUMURA_INFLAMMATORY_RESPONSE_LPS |  |  | 1.805 | 1.754 |
| REACTOME_ACTIVATED_TLR4_SIGNALLING |  |  | 1.472 |  |
| WUNDER_INFLAMMATORY_RESPONSE_AND_CHOLESTEROL_UP |  | 1.912 | 2.496 | 2.471 |
| KEGG_TOLL_LIKE_RECEPTOR_SIGNALING_PATHWAY |  |  | 1.875 | 1.778 |
| ALTEMEIER_RESPONSE_TO_LPS_WITH_MECHANICAL_VENTILATION |  | 1.875 | 2.914 | 2.774 |
| REACTOME_TOLL_RECEPTOR_CASCADES |  |  | 1.619 | 1.526 |
| REACTOME_TRIF_MEDIATED_TLR3_SIGNALING |  |  | 1.685 | 1.649 |
| ZHOU_INFLAMMATORY_RESPONSE_FIMA_UP |  |  | 1.753 | 1.535 |
| ZHOU_INFLAMMATORY_RESPONSE_LIVE_UP |  |  | 2.261 | 2.030 |
| ZHOU_INFLAMMATORY_RESPONSE_LPS_UP |  |  | 2.066 | 1.820 |
| ZHANG_INTERFERON_RESPONSE |  | 2.045 | 2.242 | 2.040 |
| REACTOME_INTERFERON_SIGNALING |  | 1.788 | 2.326 | 2.316 |
| REACTOME_INTERFERON_GAMMA_SIGNALING |  | 1.603 | 2.166 | 2.140 |
| REACTOME_IL1_SIGNALING |  |  | 1.452 |  |
| MAHAJAN_RESPONSE_TO_IL1A_UP |  | 1.642 | 1.969 | 1.589 |
| BIOCARTA_IL1R_PATHWAY |  |  | 1.958 | 1.693 |
| REACTOME_TRAF6_MEDIATED_NFKB_ACTIVATION |  | 1.698 | 1.541 | 1.511 |
| JAIN_NFKB_SIGNALING |  |  | 1.439 | 1.444 |
| REACTOME_RIP_MEDIATED_NFKB_ACTIVATION_VIA_DAI |  | 1.736 | 1.806 | 1.743 |
| HINATA_NFKB_TARGETS_KERATINOCYTE_UP |  |  | 2.405 | 2.239 |
| HINATA_NFKB_TARGETS_FIBROBLAST_UP |  |  | 2.218 | 2.182 |
| RASHI_NFKB1_TARGETS |  |  | 2.336 | 2.170 |
| MANTOVANI_NFKB_TARGETS_UP |  |  | 2.084 | 2.073 |
| REACTOME_TAK1_ACTIVATES_NFKB_BY_PHOSPHORYLATION_AND_ACTIVATION_OF_IKKS_COMPLEX |  |  | 1.631 | 1.518 |

**Supplementary Table 2.** 65-gene signature in CXCR3^+^ breast cancer cells.

| **Gene** | **Linear FC** | **P value** | **BH-adjusted P value** |
| --- | --- | --- | --- |
| *AADAC* | 2.8154 | 0.0019 | 0.0773 |
| *ABCA13* | 2.9828 | 0.0005 | 0.0368 |
| *AMIGO2* | 2.4061 | 0.0024 | 0.0553 |
| *CCDC69* | 2.3784 | 0.0020 | 0.0407 |
| *CCL20* | 2.6882 | 0.0008 | 0.0317 |
| *CCL5* | 3.3558 | 0.0002 | 0.0317 |
| *CD1D* | 3.3792 | 0.0005 | 0.0516 |
| *CD33* | 2.6329 | 0.0024 | 0.0773 |
| *CD55* | 2.8481 | 0.0007 | 0.0397 |
| *CD70* | 2.5847 | 0.0013 | 0.0429 |
| *CLCA2* | 2.9282 | 0.0012 | 0.0655 |
| *CLDN1* | 2.7511 | 0.0014 | 0.0632 |
| *CPA4* | 2.4566 | 0.0014 | 0.0358 |
| *CYTIP* | 2.3729 | 0.0026 | 0.0560 |
| *DHRS3* | 2.8679 | 0.0005 | 0.0317 |
| *EDIL3* | 2.9759 | 0.0006 | 0.0414 |
| *EHF* | 3.0035 | 0.0009 | 0.0553 |
| *EPHA4* | 2.6451 | 0.0011 | 0.0414 |
| *FAM83B* | 2.8089 | 0.0010 | 0.0498 |
| *FCAR* | 2.5491 | 0.0060 | 0.1311 |
| *FCRLA* | 2.4967 | 0.0018 | 0.0516 |
| *FYB* | 2.8154 | 0.0006 | 0.0317 |
| *GJB2* | 2.6635 | 0.0008 | 0.0324 |
| *GJB5* | 3.6553 | 0.0010 | 0.0827 |
| *GPR87* | 2.5198 | 0.0019 | 0.0553 |
| *IL1B* | 2.5257 | 0.0040 | 0.0991 |
| *IL33* | 2.8350 | 0.0011 | 0.0560 |
| *IL6* | 2.7195 | 0.0018 | 0.0719 |
| *ITGB4* | 2.4061 | 0.0023 | 0.0544 |
| *KRT14* | 2.6635 | 0.0010 | 0.0397 |
| *KRT15* | 2.3894 | 0.0109 | 0.1687 |
| *KRT17* | 2.7007 | 0.0014 | 0.0553 |
| *KRT6A* | 2.7132 | 0.0030 | 0.0973 |
| *KRT75* | 2.3620 | 0.0060 | 0.1107 |
| *LOC100505946* | 3.2868 | 0.0003 | 0.0324 |
| *LOC101927787* | 2.5315 | 0.0026 | 0.0722 |
| *LOC201651* | 2.7132 | 0.0015 | 0.0616 |
| *LPAR3* | 2.7195 | 0.0011 | 0.0482 |
| *MAB21L3* | 3.8637 | 0.0002 | 0.0398 |
| *MIR205* | 2.6268 | 0.0019 | 0.0686 |
| *MMP10* | 2.7766 | 0.0077 | 0.1687 |
| *MMP13* | 6.0210 | 0.0000 | 0.0169 |
| *MMP3* | 3.4422 | 0.0006 | 0.0627 |
| *MRGPRX3* | 3.9632 | 0.0001 | 0.0317 |
| *MT2A* | 2.5315 | 0.0028 | 0.0773 |
| *NEURL1B* | 2.9282 | 0.0022 | 0.0951 |
| *NTN4* | 2.7830 | 0.0009 | 0.0417 |
| *NUAK2* | 2.5257 | 0.0058 | 0.1249 |
| *NUP62CL* | 2.9214 | 0.0008 | 0.0498 |
| *OASL* | 3.5554 | 0.0002 | 0.0355 |
| *P2RY1* | 2.7511 | 0.0027 | 0.0951 |
| *PAK6* | 3.2944 | 0.0004 | 0.0401 |
| *PHEX* | 2.3403 | 0.0140 | 0.1865 |
| *PI3* | 3.1602 | 0.0015 | 0.0859 |
| *PKIA* | 2.4396 | 0.0028 | 0.0701 |
| *PODXL* | 2.3729 | 0.0021 | 0.0419 |
| *PPP1R14C* | 2.8547 | 0.0008 | 0.0417 |
| *PRSS3* | 2.7007 | 0.0009 | 0.0407 |
| *SDC4* | 2.4737 | 0.0022 | 0.0604 |
| *SERPINB2* | 3.5390 | 0.0011 | 0.0835 |
| *SERPINB5* | 2.5315 | 0.0022 | 0.0655 |
| *SHISA2* | 2.5491 | 0.0016 | 0.0505 |
| *SLC35F3* | 2.6208 | 0.0020 | 0.0686 |
| *SULF1* | 2.4967 | 0.0015 | 0.0414 |
| *TRPV3* | 3.2565 | 0.0013 | 0.0809 |

**Supplementary Table 3.** GO term analysis of genes enriched in CXCR3+ sorted SUM-LM1 cells (out of top 300 from PCA clustering).

| Category | Term | Fold Enrichment | BH-P value | FDR |
| --- | --- | --- | --- | --- |
| BP | **GO:0032602~chemokine production** | **22.857** | **0.000** | **0.000** |
|  | **GO:0032722~positive regulation of chemokine production** | **22.271** | **0.002** | **0.012** |
|  | **GO:0032642~regulation of chemokine production** | **19.029** | **0.001** | **0.003** |
|  | **GO:0050729~positive regulation of inflammatory response** | **11.548** | **0.005** | **0.053** |
|  | GO:0042098~T cell proliferation | 8.874 | 0.005 | 0.054 |
|  | GO:0032103~positive regulation of response to external stimulus | 7.229 | 0.002 | 0.016 |
|  | GO:0043410~positive regulation of MAPK cascade | 4.776 | 0.005 | 0.064 |
|  | GO:0030334~regulation of cell migration | 4.227 | 0.002 | 0.016 |
|  | GO:0043408~regulation of MAPK cascade | 3.985 | 0.005 | 0.069 |
|  | GO:2000145~regulation of cell motility | 3.934 | 0.004 | 0.037 |
|  | GO:0040012~regulation of locomotion | 3.770 | 0.005 | 0.059 |
|  | GO:0051270~regulation of cellular component movement | 3.610 | 0.006 | 0.094 |
|  | GO:0016477~cell migration | 3.586 | 0.000 | 0.001 |
|  | GO:0023014~signal transduction by protein phosphorylation | 3.521 | 0.005 | 0.062 |
|  | GO:0051674~localization of cell | 3.483 | 0.000 | 0.000 |
|  | GO:0048870~cell motility | 3.483 | 0.000 | 0.000 |
|  | GO:0040011~locomotion | 3.280 | 0.000 | 0.000 |
|  | GO:0006928~movement of cell or subcellular component | 2.903 | 0.000 | 0.001 |
|  | GO:0051240~positive regulation of multicellular organismal process | 2.832 | 0.005 | 0.049 |
|  | GO:0009888~tissue development | 2.748 | 0.002 | 0.009 |
|  | GO:0032268~regulation of cellular protein metabolic process | 2.304 | 0.005 | 0.047 |
|  | GO:0007166~cell surface receptor signaling pathway | 2.242 | 0.003 | 0.022 |
|  | GO:0044707~single-multicellular organism process | 1.754 | 0.000 | 0.001 |
| CC | GO:0009986~cell surface | 3.833 | 0.002 | 0.074 |
|  | GO:0005887~integral component of plasma membrane | 3.203 | 0.000 | 0.000 |
|  | GO:0005615~extracellular space | 3.191 | 0.000 | 0.003 |
|  | GO:0031226~intrinsic component of plasma membrane | 3.079 | 0.000 | 0.001 |
|  | GO:0044459~plasma membrane part | 2.305 | 0.001 | 0.022 |
|  | GO:0044421~extracellular region part | 2.053 | 0.000 | 0.006 |
|  | GO:0005576~extracellular region | 1.903 | 0.000 | 0.008 |
|  | GO:0071944~cell periphery | 1.754 | 0.001 | 0.035 |
| MF | GO:0005102~receptor binding | 2.819 | 0.005 | 0.040 |
|  | GO:0060089~molecular transducer activity | 2.687 | 0.008 | 0.028 |
|  | GO:0004872~receptor activity | 2.687 | 0.008 | 0.028 |

